# Phosphorylation Protects Oncogenic RAS from LZTR1-Mediated Degradation

**DOI:** 10.64898/2026.01.07.698128

**Authors:** Lin Zhang, Arnold Bolomsky, Omar S. Al-Odat, Callie VanWinkle, Aaliyah Battle, Papiya Chakraborty, Ronald J. Holewinski, Thorkell Andresson, Qingcai Meng, James D. Phelan, Jagan Muppidi, Ryan M. Young

**Affiliations:** Lymphoid Malignancies Branch, Center for Cancer Research, National Cancer Institute, National Institutes of Health, Bethesda, MD, USA; Protein Mass Spectrometry Group, Center for Cancer Research, National Cancer Institute, National Institutes of Health, Frederick, MD, USA; Laboratory of Cell and Molecular Biology, National Institute of Diabetes and Digestive and Kidney Diseases, National Institutes of Health, Bethesda, MD, USA

## Abstract

Oncogenic KRAS and NRAS mutations are common in hematologic malignancies, but how they signal is less well characterized than in carcinomas. To uncover novel RAS biology and potential therapeutic vulnerabilities, we employed a multi-omics screening approach in multiple myeloma to identify regulators of RAS activity. We report that the phosphatase PP1C dephosphorylates the conserved T148 residue on RAS, which in turn permits LZTR1-dependent proteasomal degradation. Notably, LZTR1 is ineffective against KRAS A146 gain-of-function mutations, which are adjacent to T148 and prevalent in hematologic cancers. Remarkably, we find that KRAS protein is four-fold less stable in hematologic versus carcinoma cells, offering a unique therapeutic opportunity targeting RAS protein stability mechanisms. The kinases PAK1 and PAK2 shield RAS from LZTR1-dependent degradation by phosphorylating T148, and targeting PAK1/2 activity improves RAS-directed therapy. Collectively, our findings reveal a novel regulatory circuit governing RAS stability that is preferentially active in blood cancers and potentially druggable.

## Main

The RAS family of small GTPases (KRAS, NRAS, and HRAS) is mutated in 30% of human cancers^1^. While principally associated with the pathogenesis of pancreatic, colon, and lung carcinomas, RAS mutations are also key drivers of many hematologic malignancies. Notably, oncogenic mutations of KRAS or NRAS are a hallmark of multiple myeloma (MM). MM is an incurable plasma cell neoplasm with over 36,000 new diagnoses this year in the US (seer.cancer.gov). Approximately 40% of newly diagnosed MM tumors harbor RAS mutations, and this figure increases to 50-70% in relapsed and refractory cases^2, 3^. As this observation implies, RAS mutations are associated with aggressive disease, and MM patients with mutant tumors have inferior outcomes^4^.

Oncogenic RAS can activate multiple signaling pathways^5^. The classical mitogen-activated protein kinase (MAPK) pathway consisting of RAF-MEK-ERK is considered a primary driver of malignant transformation in carcinomas by promoting a host of oncogenic transcription programs^6^. This signaling cascade is initiated by a complex of RAS, the adaptor SHOC2, and the serine/threonine phosphatase PP1C^7–10^, which dephosphorylates an autoinhibitory site on RAF (BRAF S365)^11^. Additionally, mutant RAS can directly activate phosphatidylinositol 3-kinase (PI3K)^12^, and this activity may be a unique feature of oncogenic RAS^13^. In contrast, our previous work demonstrated that RAS signaling in MM is qualitatively distinct from canonical signaling described in carcinomas, suggesting that RAS-dependent transformation is context-dependent. We found that pathogenic RAS signaling in MM exploits malignant plasma cell addiction to free amino acids by chronically activating mTORC1 signaling on the lysosomal surface, in parallel to its activation of the classical MAP kinase pathway^14^.

Given the dual activation of potent oncogenic pathways by RAS in MM, along with the high frequency of both KRAS and NRAS mutations in aggressive relapsed and refractory disease, effectively targeting RAS signaling represents a potential therapeutic approach for patients with tumors that have failed to respond to existing therapies. Direct targeting of KRAS and NRAS activity remains challenging, and resistance in solid tumors to approved KRAS inhibitors^15, 16^ emerges quickly^17^. A better understanding of RAS biology in hematologic cancers will help identify potential new avenues to disrupt RAS signaling.

Here, we conducted multi-omics screening to uncover novel regulators of RAS protein stability in MM. These screens and follow-up experiments reveal unique biology governing oncogenic RAS signaling specific to MM and other hematologic cancers, centered on the phosphorylation-dependent, LZTR1-mediated degradation of mutant RAS isoforms.

## Results

### Multi-omic screening identifies PPP1R2 and PP1C as regulators of KRAS protein stability

To identify novel regulators of oncogenic RAS, we developed a CRISPR screening approach using mutant KRAS^G12D^ protein expression as an endpoint (Fig. 1A; S1A). We employed KRAS-dependent XG7 and RPMI 8226 MM cell lines as model systems for RAS signaling in hematologic malignancies. Cells were engineered to ectopically express an mNeonGreen(mNG)-KRAS^G12D^–IRES–Lyt2 (mouse *Cd8a*) construct, which stoichiometrically expressed mNG-KRAS^G12D^ and Lyt2 proteins via the Internal Ribosome Entry Site (IRES).

**Figure 1.**
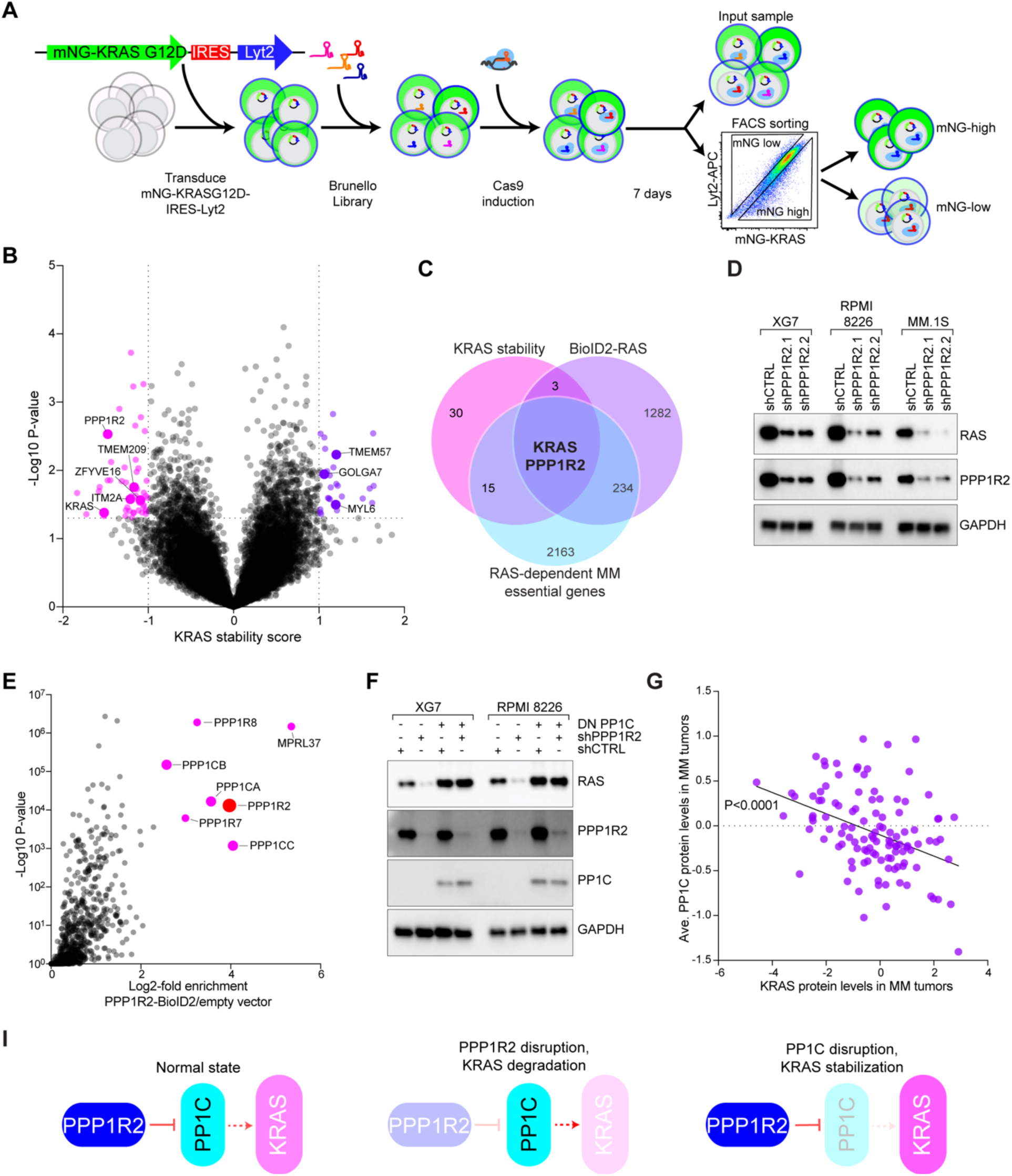
CRISPR screening reveals that PPP1R2 and PP1C control KRAS protein stability. **A**) CRISPR screening strategy to identify regulators of KRAS protein stability. **B**) Volcano plot of average KRAS stability scores (n=3). Significant hits for genes that either decrease (pink) or increase (purple) KRAS expression are indicated. Callouts represent genes identified as RAS interaction partners by mass spectrometry ^14^. **C**) Venn diagram of overlapping genes from KRAS stability screen (≤-0.8 KRAS stability score and P≤0.05), RAS BioID2 proteomics (≥1.0 log2 enrichment vs. control, from ref: ^14^), and genes essential in RAS-dependent MM cell lines (≤-1.0 CSS, from ref: ^19^). **D**) Western blot analysis of RAS, PPP1R2, and GAPDH 3 days after transduction with control shRNA (shCTRL) or PPP1R2-targeting shRNAs in XG7, RPMI 8226, and MM.1S MM lines, n=3. **E**) PPP1R2-BioID2 enrichment over empty vector in RPMI 8226 cells. **F**) Western blot analysis of RAS, PPP1R2, PP1C, and GAPDH 3 days after transduction with shCTRL, shPPP1R2.1, and/or ectopic expression of DN PP1C, n=3. **G**) Comparison of protein expression levels between KRAS (x-axis) and average PP1C (PPP1CA, PPP1CB, PPP1CC, and PPP1CC;PPP1CB) in 115 MM patient tumors. Display line is simple linear regression; R^2^=0.1593, P<0.0001. **I**) Model of PPP1R2 and PP1C regulation of KRAS protein expression. Under a normal state, PPP1R2 inhibits PP1C activity. Following PPP1R2 disruption, PP1C activity reduces KRAS levels. In contrast, PP1C disruption increases KRAS expression.

Following transduction with the Brunello single guide RNA (sgRNA) library^18^ and Cas9 induction, cells were sorted into mNG-KRAS^G12D^ high and low subpopulations. Since mNG-KRAS^G12D^ and Lyt2 expression are linked, we sorted cells for high or low mNG-KRAS^G12D^ relative to Lyt2 to control for expression differences from the construct (Fig. 1A). We calculated a “KRAS stability score”, defined as a z-score of the log2 fold change in abundance of sgRNAs for each gene between mNG-KRAS^G12D^ high and low subpopulations relative to the input sample. Data from three independent experiments (two in XG7, one in RPMI 8226) were used to generate KRAS stability scores with corresponding p-values (Fig. 1B, Table S1). The screens identified 50 genes potentially associated with decreased KRAS expression levels (≤-0.8 KRAS stability score and p≤0.05), including knockout of KRAS itself, validating the screening methodology.

These 50 genes were prioritized for further study by cross-referencing proteins proximal to mutant RAS in proteomics experiments (≥1.0 log2 enrichment vs. control)^14^ and genes essential in RAS-dependent MM lines (≤-1.0 CRISPR screen score (CSS))^19^. This analysis revealed PPP1R2 and KRAS as genes that control KRAS expression (Fig. 1C). Western blot of RAS expression following PPP1R2 knockdown confirmed that disruption of PPP1R2 expression substantially reduced RAS expression in KRAS-dependent MM cell lines XG7, RPMI 8226, and MM.1S (Fig. 1D).

PPP1R2 is a regulator of the serine/threonine phosphatase catalytic subunit PP1C^20^. PP1C controls a host of cellular processes including cell metabolism and division^21–23^. Notably, PP1C closely associates with RAS family proteins via SHOC2 to potentiate downstream MAP kinase signaling through dephosphorylation of an autoinhibitory site on RAF kinases^11^. We confirmed that SHOC2 knockdown reduced PPP1R2 and PP1C co-immunoprecipitation with KRAS^G12D^ (Fig. S1B). To characterize PPP1R2 function in MM, we tagged PPP1R2 with the promiscuous biotin ligase BioID2^24^ and identified proteins within 10-30 nm of PPP1R2 by mass spectrometry of streptavidin-purified, biotinylated proteins. As expected, PPP1R2 was closely associated with all PP1C isoforms, PPP1CA, PPP1CB, and PPP1CC, in both XG7 and RPMI 8226 MM cell lines (Fig. 1E, S1C, Table S2). Although we did not observe RAS in these PPP1R2 BioID2 experiments, we had previously found that PPP1R2 and all three PP1C isoforms were enriched in RAS BioID2 experiments performed in MM cells (Fig. S1C)^14^. PP1C genes did not significantly score in our CRISPR KRAS protein stability screens (Table S1), likely because deletion of individual isoforms would be compensated by remaining PP1C isoforms.

Depending on the context, PPP1R2 has been described as either a positive or negative regulator of PP1C activity^20^. To examine PP1C activity directly, we expressed a dominant-negative (DN) version of PP1C (PPP1CA^D95N^)^25^ in KRAS-dependent MM lines XG7 and RPMI-8226. We found that DN PP1C markedly amplified RAS expression (Fig. S1D), suggesting that PPP1R2 acts as an inhibitor of PP1C activity in this context. Concomitant knockdown of PPP1R2 and ectopic expression of DN PP1C increased RAS expression, abolishing the effect of PPP1R2 knockdown and demonstrating that PPP1R2 regulates KRAS protein stability through PP1C (Fig. 1F). Consistent with this, we found that total PP1C protein levels were inversely proportional to KRAS protein expression as measured by mass spectrometry in primary MM patient tumors^26^ (Fig. 1G; P<0.0001). These data are consistent with a model in which PP1C activity disrupts KRAS expression, and PPP1R2 acts as a negative regulator of PP1C (Fig. 1I).

### PP1C regulates KRAS phosphorylation at T148

Given that PP1C is a serine/threonine phosphatase known to be closely associated with RAS^7–9^, we hypothesized that PP1C may regulate KRAS phosphorylation. KRAS has been reported to be phosphorylated at numerous sites, including Y4, Y32, Y64, S89, Y137, T144, T148, and S181, which can regulate its activity and stability^27^. To identify potential serine/threonine phosphorylation sites that control KRAS stability in MM, we generated eight S/T to A mutants on the mNG-KRAS^G12D^-IRES-Lyt2 backbone based on previously characterized phosphorylation sites^28^ (Fig. 2A). Using these constructs, we performed FACS analysis of mNG-KRAS expression in MM cell lines expressing similar expression levels as judged by Lyt2 surface staining (Fig. S2A). Among the eight sites, disruption of T148 consistently reduced mNG-KRAS^G12D^ protein expression as measured by FACS (Fig. 2B-C) or western blot (Fig. S2B).

**Figure 2.**
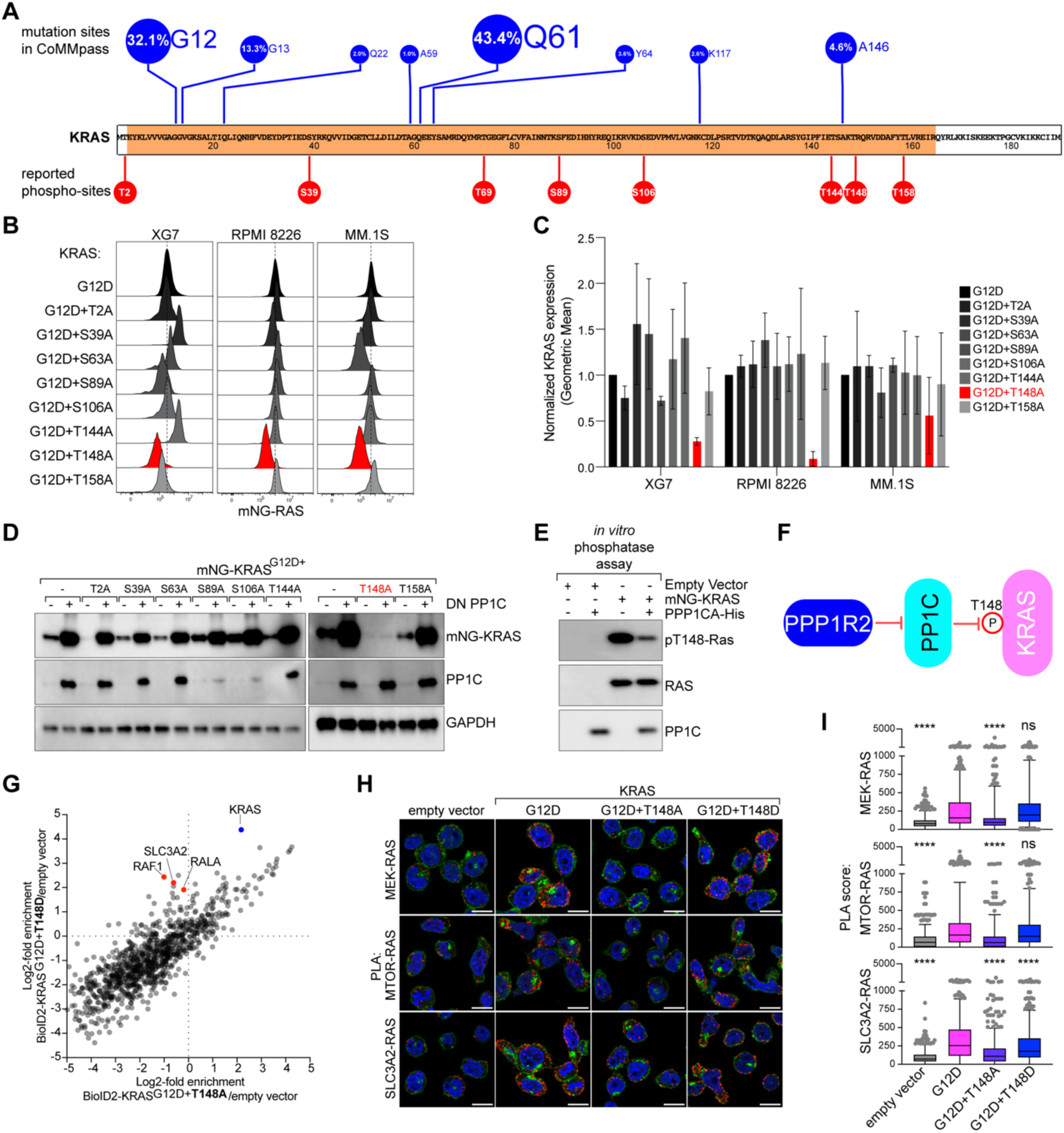
PP1C dephosphorylates KRAS T148. **A**) Schematic of KRAS oncogenic hotspot mutations and putative serine/threonine phosphorylation sites. **B**) Representative FACS data for mNG-KRAS^G12D^ constructs harboring indicated S/T to A mutations XG7, RPMI 8226, and MM.1S MM cells, n=3. **C**) Average normalized mNG-KRAS^G12D^ phospho-mutant FACS data. Error bars depict standard deviation, n=3. **D**) Western blot analysis for mNG-KRAS, PP1C, and GAPDH in XG7 cells expressing indicated mNG-KRAS mutants and either empty vector (-) or DN PP1C (+), n=3. **E**) Western blot analysis of in vitro phosphatase assays performed on mNG-KRAS^G12D^ immunoprecipitated from XG7 cells, n=4. **F**) Model depicting PP1C directly dephosphorylating KRAS T148. **G**) Enrichment of KRAS^G12D+T148A^ (x-axis) vs. KRAS^G12D+T148D^ (y-axis) BioID2 constructs. **H**) Representative images from proximity ligation assay (PLA) (red) of RAS and indicated downstream effectors in XG7 cells. DAPI (blue), wheat germ agglutinin (WGA; green). Scale bar is 10 μm. **I**) PLA score of indicated PLA pairs normalized to empty vector condition, **** P<0.0001, n=3.

To examine whether PP1C regulates phosphorylation at T148, we expressed DN PP1C in cells expressing the eight KRAS^G12D^ phospho-mutants. Expression of KRAS^G12D^ harboring the T148A mutation did not increase with DN PP1C expression, while the seven other phospho-mutants and the KRAS^G12D^ control each showed increased KRAS expression (Fig. 2D).

Similarly, KRAS^G12D+T148A^ expression was not further reduced by PPP1R2 knockdown, whereas all other KRAS^G12D^ phospho-mutants had less expression (Fig. S2C). Additionally, KRAS^G12D+T148A^ could not rescue XG7 survival as effectively as KRAS^G12D^ following CRISPR-mediated knockout of endogenous mutant KRAS (Fig. S2D). We also generated a T148D phospho-mimetic mutation in KRAS^G12D^ to model constitutive KRAS T148 phosphorylation.

KRAS^G12D+T148D^ showed elevated protein expression compared to KRAS^G12D^, and its expression was insensitive to modulation of PP1C activity through either DN PP1C overexpression or PPP1R2 knockdown (Fig. S2E-F). These data corroborate that PPP1R2 and PP1C regulate KRAS phosphorylation at T148.

We next developed a phospho-specific antibody to directly examine endogenous T148 phosphorylation (Fig. S2G). KRAS T148 phosphorylation increased upon DN PP1C expression in KRAS-dependent XG7, RPMI 8226, and MM.1S cells (Fig. S2H-I), demonstrating that PP1C regulates phosphorylation of endogenous mutant KRAS. To determine whether PP1C directly dephosphorylated KRAS, we performed an *in vitro* phosphatase assay. XG7 cells expressing mNG- KRAS^G12D^ were lysed in SDS and boiled to disrupt protein-protein interactions.

KRAS^G12D^ was then immunoprecipitated with anti-mNG-beads and combined with recombinant HIS-tagged PPP1CA for 1 hour. Western blot analysis using anti-phospho-T148-RAS demonstrated that PP1C directly dephosphorylated KRAS T148 (Fig. 2E-F).

To assess the functional impact of KRAS T148 phosphorylation, we cloned BioID2-tagged versions of KRAS^G12D^ with T148A and T148D substitutions and compared interaction partners between unphosphorylated and phosphorylated models of KRAS^G12D^ by mass spectrometry.

KRAS^G12D+T148D^ demonstrated significantly higher expression and had increased association with known RAS signaling effectors in MM, including RAF1, RALA, and SLC3A2, relative to KRAS^G12D+T148A^ (Fig. 2G, Table S3)^14^. We performed proximity ligations assays (PLAs) to confirm protein-protein associations between expressed KRAS phospho-mutants and downstream effectors MEK, MTOR, and SLC3A2. PLA is an antibody-based method that can detect protein-protein associations within ∼40 nm visualized as red fluorescent puncta ^29^.

KRAS^G12D+T148A^ had significantly lower associations with these key signaling proteins versus KRAS^G12D^ or KRAS^G12D+T148D^ (Fig. 2H-I).

### T148 phosphorylation protects KRAS from LZTR1-mediated ubiquitination

Since PPP1R2/PP1C regulates KRAS protein stability through T148 phosphorylation, we hypothesized that this site may control KRAS ubiquitination and subsequent proteasomal degradation. Consistent with this, inhibiting proteasomal degradation with MG-132 treatment increased KRAS^G12D+T148A^ protein expression (Fig. S3A). Therefore, we directly examined the ubiquitination status of KRAS^G12D^ and its T148A and T148D phospho-mutants expressed in XG7 and RPMI 8226 MM cells. Compared to KRAS^G12D^, the non-phosphorylatable KRAS^G12D+T148A^ had increased ubiquitin binding, while the phospho-mimicking KRAS^G12D+T148D^ demonstrated reduced ubiquitin binding (Fig. 3A). Likewise, modulating PPP1R2 and PP1C expression and activity modified KRAS ubiquitination (Fig. S3B-C), confirming that phosphorylation at T148 prevents RAS degradation.

**Figure 3.**
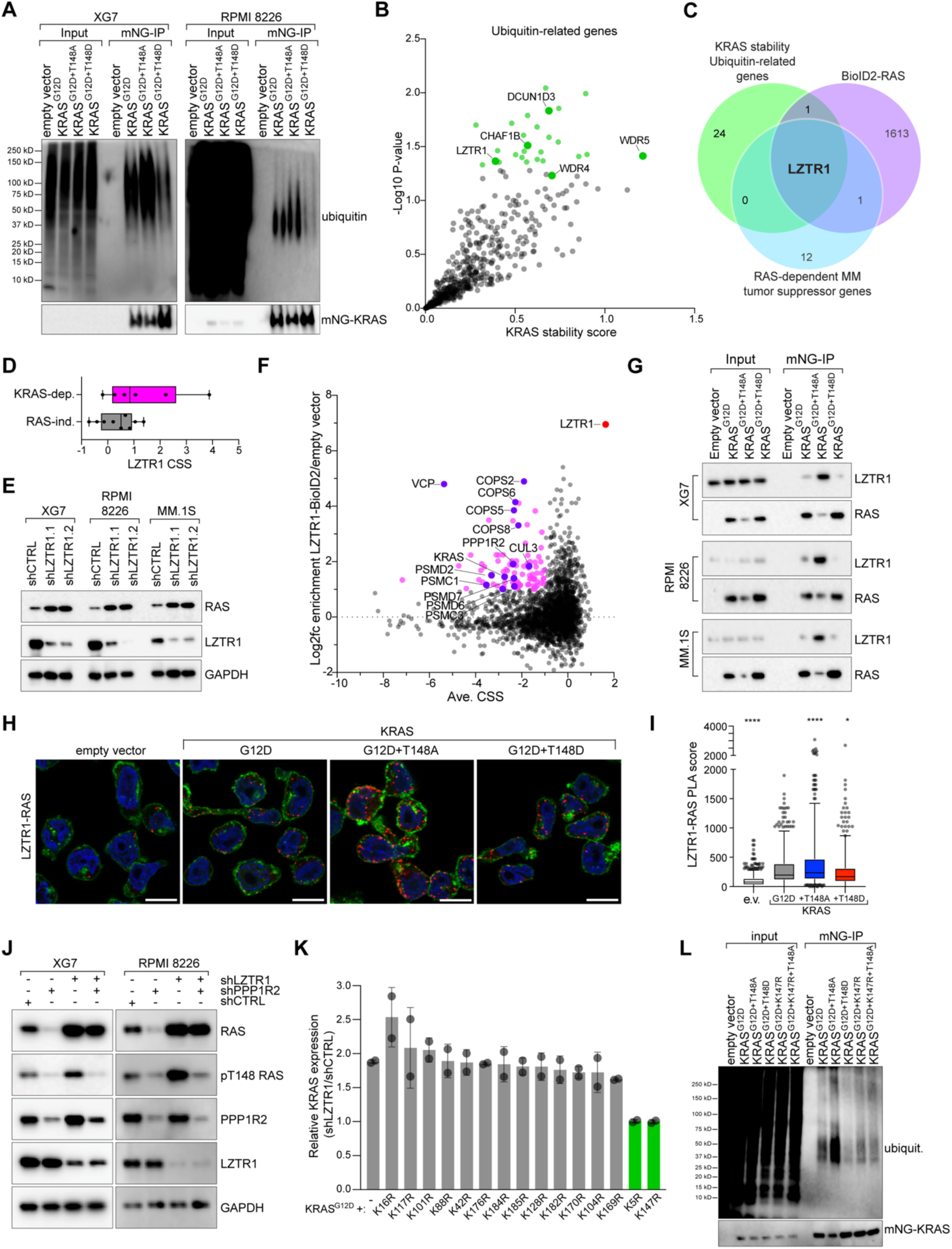
T148 phosphorylation protects KRAS from LZTR1-mediated degradation. **A**) IP-western blot analysis of ubiquitin binding to indicated ectopically expressed KRAS constructs. **B**) Scatter plot of ubiquitin-related genes from the KRAS stability screens in Fig. 1A-B. **C**) Venn diagram of ubiquitin-related genes with increased KRAS expression (P≤0.05), KRAS interaction partners from BioID2 studies (≥1.0 log2 fold enrichment, ref: ^14^), and tumor suppressor genes identified in RAS-dependent MM lines (CSS≥0.5, ^19^). **D**) LZTR1 CSS ^19^ for KRAS-dependent (pink) vs. RAS-independent (gray) MM lines. **E**) Western blot analysis of RAS, LZTR1, and GAPDH following expression of shCTRL or 2 shRNAs targeting LZTR1 in XG7, RPMI 8226, and MM.1S MM lines. **F**) The essential interactome of LZTR1. Average CSS for RPMI 8226 and XG7 (x-axis) plotted against average enrichment for proteins from LZTR1-BioID2 experiments in the same lines (y-axis). Pink shading denotes proteins with ≥1.0 log2 fold enrichment and CSS ≤1; purple are callouts discussed in text. **G**) Co-IP of indicated KRAS^G12D^ phospho-mutants with LZTR1 in XG7, RPMI 8226, and MM.1S MM cells. **H**) Representative images from PLA (red) of RAS and LZTR1 in XG7 cells expressing indicated KRAS mutants. DAPI (blue), wheat germ agglutinin (WGA; green). Scale bar is 10 μm. **I**) PLA score of LZTR1-RAS normalized to empty vector condition, **** P<0.0001, * P<0.01, n=3. **J**) Western blot analysis of RAS, pT148-RAS, PPP1R2, LZTR1, and GAPDH following transduction with control, PPP1R2, and/or LZTR1 shRNAs. **K**) Bar graph of relative changes in mNG-KRAS expression measured by FACS for indicated mutants in shLZTR1 vs. shCTRL treated cells. **L**) Immunoblot of ubiquitin binding following mNG-KRAS pulldown in cells transduced harboring G12D, T148A, T148D, K147R or K147R+T148A mutations with ubiquitin in XG7 cells, n=2.

We next sought to identify potential ubiquitin ligases responsible for T148-dependent KRAS ubiquitination. We focused on ubiquitin-related genes^30^ that significantly increased KRAS expression in the stability screens (p≤0.05) (Fig. 3B). Comparing these genes with KRAS interaction partners (≥1.0 log2 fold enrichment)^14^ and tumor suppressor genes in RAS-dependent MM lines (CSS≥0.5)^19^ identified LZTR1 (Fig. 3C). LZTR1 is an E3 ubiquitin ligase previously reported to promote RAS degradation^31, 32^, along with RAS-related proteins such as RIT1^33^.

Accordingly, LZTR1 has a tumor suppressor phenotype following CRISPR-mediated knockout in KRAS-dependent, but not RAS-independent, MM lines (Fig. 3D), and LZTR1 knockdown amplified KRAS protein expression in KRAS-dependent MM lines (Fig. 3E) via decreased K48-linked ubiquitination of KRAS (Fig. S3D). Moreover, LZTR1 knockdown significantly increased the half-life of RAS protein in XG7 and MM.1S cells treated with cycloheximide (Fig. S3E-F).

To gain insight into the function of LZTR1 in MM cells, we ectopically expressed an LZTR1-BioID2 construct in KRAS-dependent RPMI 8226 and XG7 cells. We observed strong interactions between LZTR1 and CUL3, various proteasomal subunits (PSMC1, PSMD2, PSMD7, PSMD6, PSMC3), and the COP9 signalosome (COPS2, COPS6, COPS5, COPS8), consistent with its function as an adaptor for the CUL3-RBX1 E3 ubiquitin ligase complex (Fig. 3F; Table S4). Gene Ontology^34^ enrichment of proteins associated with LZTR1 (≥1.0 log2 fold enrichment) identified strong concordance with regulation of organelle enrichment and vesicle-mediated transport (Fig. S3G). Moreover, we found that LZTR1 associated with both KRAS and PPP1R2 proteins and, to a lesser extent, PP1C family members, suggesting that LZTR1 may cooperate with phosphorylation-dependent regulation of KRAS. Indeed, LZTR1 preferentially associated with the KRAS^G12D+T148A^ phospho-mutant by co-immunoprecipitation (Fig. 3G) and PLA (Fig. 3H-I), suggesting KRAS T148 controls LZTR1-mediated degradation.

We next disrupted PPP1R2 and LZTR1 expression in KRAS-dependent XG7 and RPMI 8226 MM lines. PPP1R2 knockdown, which increases PP1C activity, reduced both total and phospho-T148 KRAS levels, whereas LZTR1 knockdown increased both total and phospho-T148 KRAS (Fig. 3J). Simultaneous knockdown of LZTR1 and PPP1R2 increased total RAS while reducing phospho-T148 RAS, demonstrating that PPP1R2-dependent loss of KRAS expression is mediated by LZTR1.

To understand how LZTR1 regulates KRAS stability in MM, we mutated lysine residues on mNG-KRAS^G12D^ to arginine to prohibit ubiquitination while maintaining the positive charge (Fig. S3H). Constructs were expressed in XG7 cells and KRAS protein stability was monitored by FACS analysis following LZTR1 knockdown (Fig. 3K). Among the 15 lysine residues we tested, only mutation of KRAS K5 and K147 rendered KRAS expression insensitive to LZTR1 knockdown. Notably, K147 is adjacent to the T148 phosphorylation site on KRAS. Therefore, we examined ubiquitin binding to ectopically expressed KRAS^G12D^ mutants harboring K147 and T148A mutations (Fig. 3L). As before, KRAS^G12D+T148A^ had substantially more ubiquitin binding the KRAS^G12D^ alone, whereas KRAS^G12D+T148D^ showed reduced binding. Interestingly, KRAS^G12D+K147R^ had less ubiquitin binding than KRAS^G12D^ and further addition of T148A failed to increase ubiquitin binding, suggesting that T148 phosphorylation controls K147 ubiquitination. Recently, a crystal structure of LZTR1 bound to GDP-KRAS was solved^35^.

Modeling of GDP-bound KRAS^G12D^ with T148 phospho-mutants with LZTR1 and CUL3^36^ with AlphaFold^37^ corroborates that a negative charge at T148 disrupts LZTR1 binding proximal to K147 (yellow) on KRAS (Fig. S3I). Taken together, these data support a model in which PP1C/PPP1R2-dependent regulation of KRAS T148 phosphorylation protects KRAS from LZTR1-mediated ubiquitination of K147 (Fig. S3J).

### KRAS A146 mutations are resistant to LZTR1-mediated degradation

While we found that K147 can negatively regulate KRAS expression, there are no reported KRAS K147 mutations among MM tumors profiled in the Multiple Myeloma Research Foundation (MMRF) CoMMpass study^38^. However, oncogenic hotspot mutations at position A146 are common. KRAS A146 mutations (A146T, A146V, A146P) disrupt guanine nucleotide binding to promote GDP for GTP exchange^39^ and are found in MM and a handful of other cancers, including chronic myelogenous leukemia (CML), glioblastoma (GBM), (acute lymphoblastic leukemia (ALL), and acute myeloid leukemia (AML) (Fig. 4A; ref: ^1^).

**Fig. 4.**
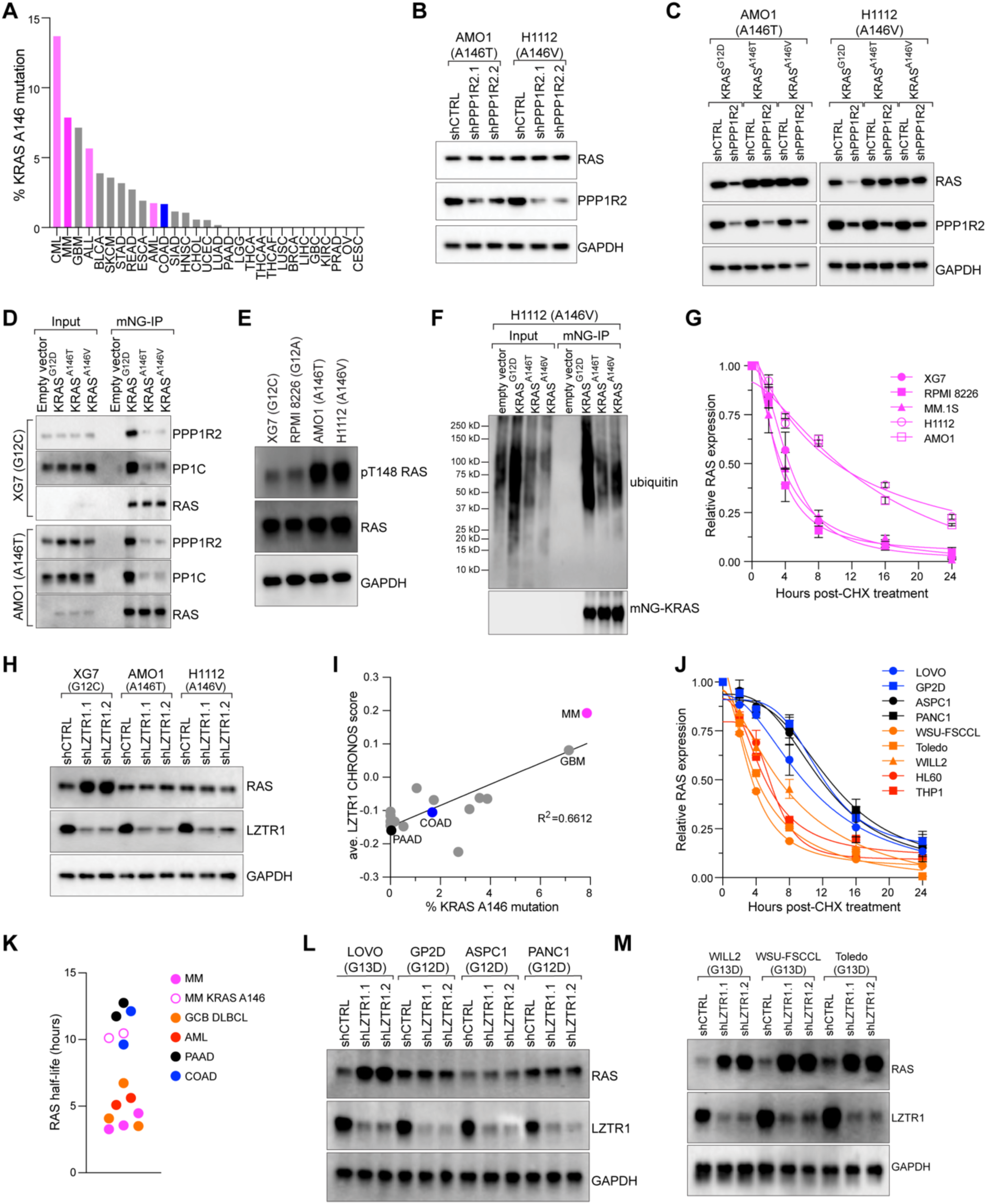
A146 mutations protect KRAS from LZTR1-mediated degradation. **A**) Bar graph of KRAS A146 mutations as a percentage of all KRAS mutations across various tumor types. Hematologic malignancies are highlighted in pink. Adapted from ref: ^1^. **B,C**) Western blot analysis of RAS, PPP1R2, and GAPDH following transduction with control or PPP1R2 shRNAs (**B**) or indicated ectopically expressed RAS mutants (**C**), n=3. **D**) Co-IP of indicated KRAS mutants with PPP1R2 or PP1C in XG7 and AMO1 MM cells, n=3. **E**) Western blot analysis of pT148 RAS phosphorylation of endogenous RAS in XG7, RPMI 8226, AMO1, and H1112 MM lines, n=3. **F**) Western blot analysis of ubiquitin binding following mNG-KRAS pulldown in H1112 cells harboring G12D, A146T, or A146V, n=3. **G**) Quantification of immunoblots of RAS expression normalized to GAPDH from indicated MM lines following a time course of treatment with 10 nM cycloheximide (CHX) for the indicated timepoints (n=2; error bars depict standard deviation; representative blots in Fig. S4G). **H**) Western blot analysis of RAS, LZTR1, and GAPDH following transduction with control or LZTR1 shRNAs in XG7, AMO1, and H1112 cells, n=3. **I**) Scatter plot showing the percentage of KRAS A146 mutations among all KRAS mutations from Fig. 4A (x-axis) versus the LZTR1 CHRONOS score from DepMap. **J**) Quantification of immunoblots of RAS expression normalized to GAPDH from indicated cell lines (COAD, blue; PAAD, black; GCB DLBCL, orange; AML, red) following a time course of treatment with 10 nM cycloheximide (CHX) for the indicated timepoints (n=2; error bars depict standard deviation; representative blots in Fig. S4G). **K**) Dot plot showing the RAS half-life in indicated cell lines (MM, pink; MM KRAS A146, pink hollow circle; COAD, blue; PAAD, black; GCB DLBCL, orange; AML, red). **L,M**) Immunoblot analysis of RAS expression following transduction with control shRNA or LZTR1 shRNAs in adenocarcinoma lines (**L**) and GCB DLBCL lines (**M**), n=3.

Given that the A146 residue is adjacent to K147 and T148, we hypothesized that KRAS A146 mutations might confer resistance to PPP1R2/PP1C/LZTR1-mediated degradation, in addition to promoting GTP exchange. To investigate this, we used AMO1 and H1112 MM cells that endogenously express KRAS^A146T^ and KRAS^A146V^, respectively. In these cells, PPP1R2 knockdown or DN PP1C expression had no effect on RAS expression in these cells (Fig. 4B, S4A). To rule out cell-intrinsic factors, we stably expressed KRAS^G12D^, KRAS^A146T^, or KRAS^A146V^ in AMO1 and H1112 cells. PPP1R2 knockdown (Fig. 4C) or DN PP1C expression (Fig. S4B) only affected KRAS^G12D^ expression, suggesting that KRAS A146 mutations are insensitive to PPP1R2/PP1C regulation. Correspondingly, we observed co-immunoprecipitation of PPP1R2 and PP1C with KRAS^G12D^, but not with KRAS^A146T^ and KRAS^A146V^ in XG7 or AMO1 MM lines (Fig. 4D). This indicates that PP1C cannot bind to, and therefore cannot dephosphorylate, KRAS harboring A146 mutations. After first confirming the anti-pT148 KRAS antibody could specifically detect T148 phosphorylation on KRAS^A146T^ (Fig. S4C), we found that AMO1 and H1112 MM cells harboring KRAS A146 mutations had higher basal levels of KRAS T148 phosphorylation relative to XG7 and RPMI 8226 bearing KRAS G12 mutations (Fig. 4E).

Our results indicate that KRAS A146 mutations confer resistance to LZTR1-mediated ubiquitination and degradation. Notably, MM patients harboring KRAS A146 mutations had significantly higher levels of LZTR1 transcripts than patients with other oncogenic KRAS or NRAS mutations, supporting the notion that these mutants are less sensitive to LZTR1 expression. Accordingly, we found that KRAS^A146T^ and KRAS^A146V^ mutations were less ubiquitinated than KRAS^G12D^ when ectopically expressed in H1112 cells (Fig. 4F).

Correspondingly, we observed increased RAS protein stability in AMO1 and H1112 cells harboring KRAS A146 mutations versus XG7, RPMI 8226, and MM.1S cells expressing KRAS G12 mutations.

These data suggest that LZTR1 may be less active against A146 mutations, and we found that LZTR1 knockdown had no effect on RAS expression levels in AMO1 and H1112 MM cells (Fig. 4H). However, we observed increased RAS levels following LZTR1 knockdown in control XG7 cells expressing KRAS^G12C^ (Fig. 4H). The addition of K147R and T148 mutations onto an mNG-KRAS^A146T^ backbone did not further alter KRAS protein expression (Fig. S4D). Together, these data support that A146 mutations confer resistance to LZTR1-mediated degradation, which is a distinct oncogenic mechanism from their promotion of GTP binding.

Interestingly, data from the DepMap CRISPR dependency database indicates that LZTR1 preferentially functions as a tumor suppressor in KRAS-dependent MM versus in KRAS-driven adenocarcinomas (Fig. S4E)^40^. We also found a significant correlation (R^2^=0.6612; p<0.0001) between the frequency of A146 mutations in tumor types and the magnitude of LZTR1’s tumor suppressor effect (Fig. 4I). MM (pink) and glioblastoma (GBM) had both the highest frequency of KRAS A146 mutations while LZTR1 exhibited the strongest tumor suppressor effects.

Conversely, most adenocarcinomas, such as colon (COAD, blue) and pancreatic adenocarcinomas (PAAD, black) had a low prevalence of KRAS A146 mutations and were not responsive to LZTR1 knockout in DepMap (Fig. 4I).

These data suggest that LZTR1 may not regulate KRAS expression in many adenocarcinomas to the same extent as in MM, consistent with the reported long half-life of KRAS^G12C^ in epithelial tumor lines (∼22 hours)^41^. Above, we found that MM cells harboring KRAS mutations at the G12 codon had a substantially shorter half-lives of around 3-4 hours. To better compare RAS protein stability between different tumor types, we used cycloheximide to track RAS protein levels in diverse cell lines which all harbored mutant KRAS, including adenocarcinoma lines from colon (COAD; LOVO and GP2D; blue) and pancreas (PAAD; ASPC1 and PANC1; black), as well as additional blood cancer lines from germinal center B cell diffuse large B cell lymphoma (GCB DLBCL) (WSU-FSCCL, Toledo, and WILL2; orange) and AML (HL60 and THP1; red). While RAS expression rapidly declined in MM cells within 3-4 hours (Fig. 4G, K), RAS remained stable for up to 12 hours in tested adenocarcinoma lines (Fig. 4J-K, S4F).

Interestingly, RAS protein was had a relatively short half-life of 4-6 hours in both GCB DLBCL and AML cell lines (Fig. 4J-K).

We next directly examined the roles of LZTR1 and PPP1R2/PP1C in regulating KRAS stability across the tested adenocarcinoma lines, as well as in the pancreatic cancer line Mia PaCa-2.

LZTR1 knockdown increased RAS protein expression in LOVO but had no effect on the other cell lines (Fig. 4L, S4G), consistent with our cycloheximide data demonstrating that LOVO had an intermediate RAS stability between the adenocarcinoma and hematologic cancer lines (Fig. 4J). Likewise, changes in RAS expression following PPP1R2 knockdown or ectopic expression of DN PP1C were observed only in LOVO, but not GP2D, ASPC1, PANC1, and Mia PaCa-2 (Fig. S4H-I).

The above data suggests that PPP1R2/PP1C and LZTR1 regulation of KRAS expression may be a feature of hematologic malignancies. To confirm this, we used cell line models of GCB DLBCL. Although KRAS mutations are rare in in GCB DLBCL, mutations targeting A146 constitute over 10% of total KRAS mutations in this cancer type^42, 43^. Consistent with our findings in MM, we observed that LZTR1 knockdown (Fig. 4M) and ectopic expression of DN PP1C (Fig. S4J) increased RAS expression, while PPP1R2 knockdown (Fig. S4K) reduced RAS in the GCB DLBCL cell lines WILL2, WSU FSCCL, and Toledo, which all express mutant KRAS^G13D^. Together, these data suggest that RAS stability is differentially regulated in hematologic cancers versus adenocarcinomas and further support the notion that PPP1R2/PP1C/LZTR1-dependent regulation of KRAS stability is a feature of hematologic cancers.

### PP1C modulates NRAS T148 phosphorylation and LZTR1-dependent degradation

MM harbors both oncogenic activating mutations of KRAS and NRAS (Fig. S5A)^1^, and while ALL and AML also harbor KRAS A146 mutations, these leukemias preferentially express mutant forms of NRAS^44^. The amino acid sequence surrounding T148 is highly conserved across human KRAS, NRAS, and HRAS (Fig. 5A), suggesting that PPP1R2/PP1C could also regulate NRAS expression via T148 phosphorylation. We found that PPP1R2 knockdown reduced NRAS expression in the NRAS-dependent MM lines SKMM1, INA6, and NCI H929, as well as the HL60 and THP1 AML lines (Fig. 5B-C). Conversely, expression of DN PP1C increased NRAS expression in these lines (Fig. S5B-C).

**Figure 5.**
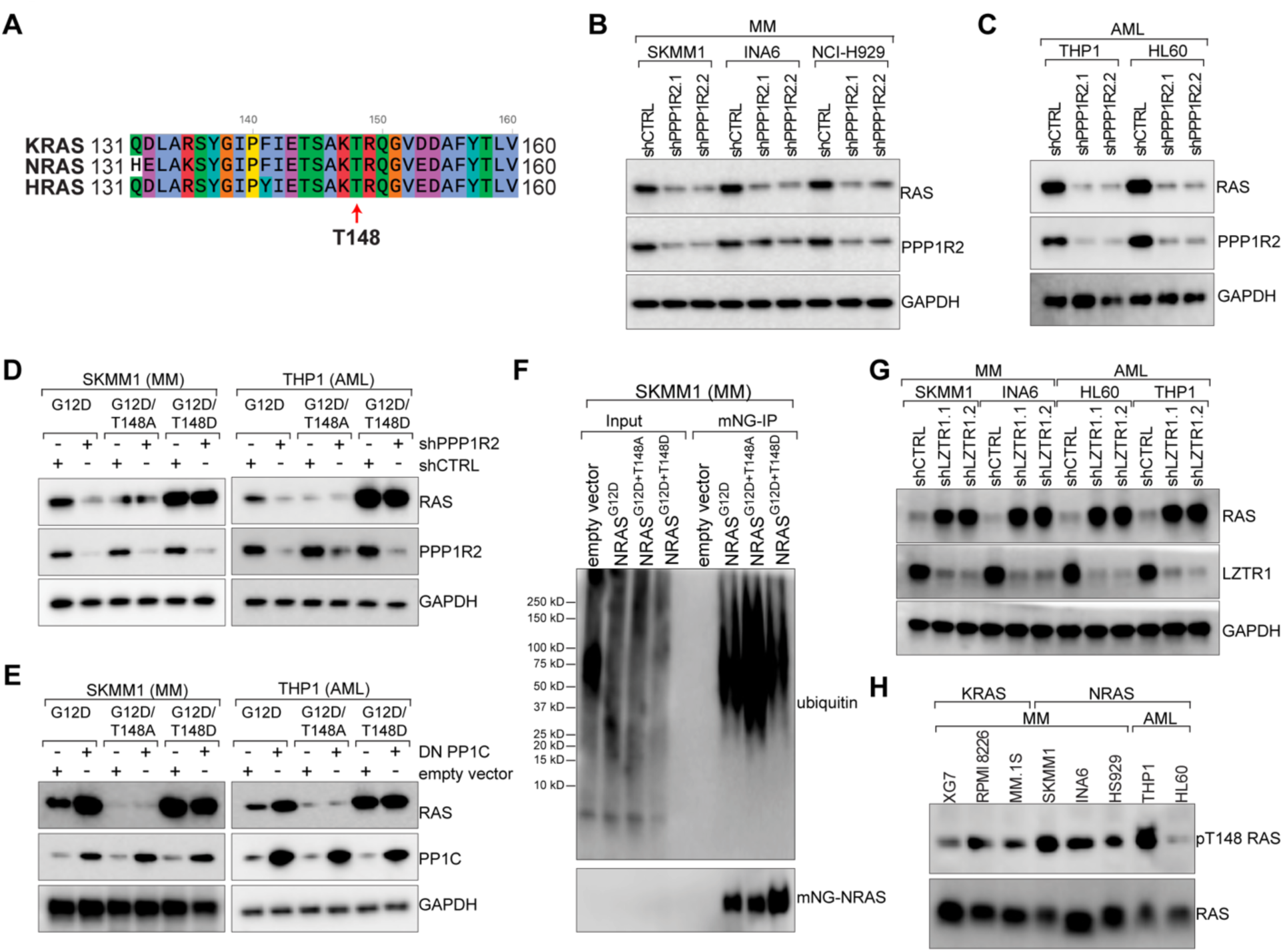
PP1C regulates NRAS T148 phosphorylation and LZTR1-dependent degradation. **A**) Sequence alignment of human KRAS (KRASA), NRAS, and HRAS surrounding T148. **B**) Western blot analysis of RAS, PPP1R2, and GAPDH following transduction with control shRNA or shRNAs targeting PPP1R2 in SKMM1, INA6, and NCI-H929 MM lines, n=3. **C**) Western blot analysis of RAS, PPP1R2, and GAPDH following transduction with control shRNA or shRNAs targeting PPP1R2 in THP1 and HL60 AML lines n=3. **D**) Ectopic expression of indicated NRAS phospho-mutants and western blot analysis of RAS expression following PPP1R2 knockdown, n=3. **E**) Ectopic expression of indicated NRAS phospho-mutants and Western blot analysis of RAS expression following DN PP1C expression in SKMM1 MM cells and THP1 AML cells, n=3. **F**) IP-western analysis of NRAS ubiquitin binding to indicated ectopically expressed NRAS mutants in SKMM1 MM cells, n=3. **G**) Western blot analysis of RAS, LZTR1, and GAPDH in SKMM1, INA6, HL60, and THP1 cells following treatment with control shRNA or shRNAs targeting LZTR1, n=3. **H**) Evaluation of pT148 RAS phosphorylation in the indicated MM and AML cell lines.

These data suggested that T148 phosphorylation could also regulate NRAS protein stability. Ectopic expression of mNG-NRAS^G12D^ constructs harboring T148 phospho-mutants showed that T148A mutation reduced NRAS expression levels as measured by FACS in SKMM1 and INA6 MM lines, and in THP1 AML cell line (Fig. S5D). Western blot analysis revealed that while PPP1R2 knockdown reduced expression NRAS^G12D^, it had no effect on NRAS phospho-mutants T148A and T148D (Fig. 5D). Similarly, overexpression of DN PP1C only increased expression of NRAS^G12D^, while the T148 phospho-mutant was unaffected in SKMM1 MM cells and THP1 AML cells (Fig. 5E).

Consistent with our findings for KRAS, we found that ectopically expressed NRAS phospho-mutants modulated ubiquitin binding, with increased ubiquitin binding observed for NRAS^G12D+T148A^ in both the SKMM1 MM line and THP1 AML cells (Fig. 5F, S5E). Moreover, NRAS protein levels increased following LZTR1 knockdown in MM and AML lines (Fig. 5G), indicating that LZTR1-mediated NRAS degradation is also regulated by T148 phosphorylation. As expected, our anti-phospho-T148 KRAS antibody also recognized T148 phosphorylation on NRAS at levels comparable to that found on KRAS in MM (Fig. 5H).

### PAK1 and PAK2 phosphorylate RAS T148

We determined that PP1C dephosphorylates KRAS and NRAS at T148, which in turn regulates LZTR1-mediated RAS degradation. This mechanism suggests the existence of a reciprocal kinase that phosphorylates T148 (Fig. 6A). GSK3B has been reported to phosphorylate RAS at T148 and T144 to increase RAS protein expression^45^, and we also found that MM cells treated with the GSK3B inhibitor IM-12^46^ modestly increased RAS expression by immunoblot (Fig. S6A).

**Figure 6.**
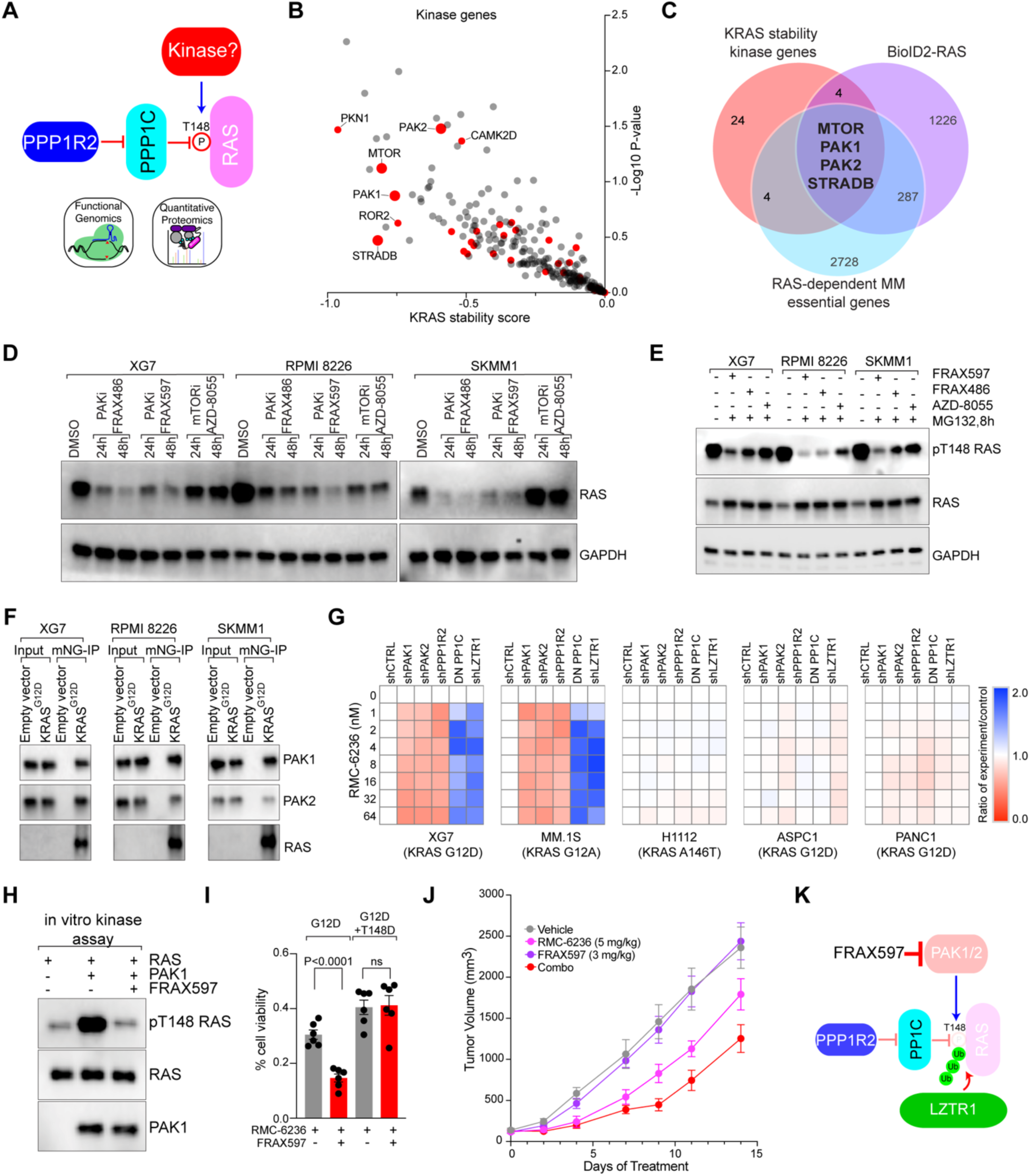
Identification of kinases targeting RAS T148. **A**) Schematic for a putative kinase phosphorylating RAS T148. **B**) Scatter plot of kinase-related genes from the KRAS stability screens in Fig. 1A-B. **C**) Venn diagram of kinase genes with decreased KRAS expression (≥0.5 KRAS stability score), RAS interaction partners from BioID2 studies (≥0.8 log2 fold enrichment, ^14^), and essential genes identified in RAS-dependent MM lines (CSS<-1.0, ^19^). **D**) Western blot analysis of RAS expression following treatment with 100 nM of the indicated drugs for 24 and 48 hours in XG7, RPMI 8226, and SKMM1 MM cells, n=5. **E**) Evaluation of pT148 RAS phosphorylation in cells treated with MG-132 for 8 hours before lysis and the indicated drugs for 48 hours, n=3. **F**) Co-IP of PAK1 and PAK2 with ectopically expressed mNG-KRAS^G12D^ in the indicated MM lines, n=2. **G**). Indicated cells expressing listed shRNAs or DN PP1C were treated with a dose titration of RMC-6236. Data shown is the ratio of experiment (listed shRNA or DN PP1C) versus shCTRL. Values shown are the average of 2 experiments. **H**) in vitro kinase assay of recombinant PAK1 on recombinant KRAS as a substrate. Western blot analysis of phospho-T148 RAS was used as a readout. Inhibition with FRAX597 was used to demonstrate specificity, n=4. **I**) Cell viability in XG7 cells ectopically expressing KRAS^G12D^ or KRAS^G12D+T148D^ and treated with indicated drugs: either 100 nM FRAX527 and/or 8 nM RMC-6236. **J**) Tumor volume for MM.1S xenografts treated with vehicle (gray), 3 mg/kg FRAX597 (purple), 5 mg/kg RMC-6236 (pink), or the combination of both drugs (red). **K**) Inhibition with FRAX597 primes RAS for degradation.

However, we sought a T148-specific kinase whose inhibition would reduce RAS expression, presenting a potential vulnerability that might be exploited therapeutically to target RAS-dependent hematologic malignancies. To identify candidate kinases, we examined kinase genes (www.kinhub.org) that reduced KRAS expression in the KRAS stability screens (Fig. 6B).

Comparing these genes (≤-0.5 KRAS stability score) with RAS interaction partners (≥0.8 log2 enrichment) and essential genes in RAS-dependent MM (CSS ≤-1.0) identified four potential kinases: MTOR, PAK1, PAK2, and STRADB (Fig. 6C).

MTOR and PAK1/2 have multiple inhibitors small molecule inhibitors that might affect RAS phosphorylation and protein stability, whereas STRADB is a pseudo-kinase and has no kinase activity or inhibitors. Inhibiting PAK1/2 with FRAX597 and FRAX486 or MTOR with AZD-8055 reduced RAS expression in MM cell lines within 24 to 48 hours, as demonstrated by immunoblot (Fig. 6D) and FACS analysis on cells ectopically expressing mNG-KRAS^G12D^ (Fig. S6B-C). Since MTOR inhibition quenches overall protein synthesis, thereby non-specifically reducing RAS, we next examined if these kinase inhibitors specifically reduced RAS T148 phosphorylation. We found that both PAK1/2 inhibitors robustly decreased RAS T148 phosphorylation in MM cells treated with MG-132, whereas the MTOR inhibitor AZD-8055 had a more muted effect on T148 phosphorylation (Fig. 6E). Accordingly, FRAX597 treatment could not reduce KRAS^G12D^ protein expression when T148 phospho-mutants were co-expressed, while AZD-8055 reduced RAS expression regardless (Fig. S6D).

These data suggest that PAK1 and PAK2 regulate RAS expression by directly phosphorylating T148. Consistent with this, both PAK1 and PAK2 co-immunoprecipitated with ectopically expressed RAS (Fig. 6F). Moreover, PAK1/2 inhibition did not reduce RAS protein expression in AMO1 and H1112 MM cells harboring KRAS A146 mutations (Fig. S6E-G), in agreement with our above data demonstrating that A146 mutations protect RAS from LZTR1-mediated degradation regardless T148 phosphorylation status. PAK1/2 inhibition also moderated the action of LZTR1 knockdown on KRAS protein levels in XG7, RPMI 8226, and SKMM1 MM lines (Fig. S6H), demonstrating that PAK1/2 modulates RAS expression in an LZTR1-dependent manner.

Treatment with FRAX597 alone was not selectively toxic to RAS-dependent MM (Fig. S6I). Although PAK1 knockout was selectively toxic to RAS-dependent MM in CRISPR screens, PAK2 proved essential gene regardless of RAS dependency (Fig. S6J). To determine whether PAK1 or PAK2 preferentially regulated RAS stability, we generated MM (XG7, MM.1S, H1112) and PAAD (ASPC1, PANC1) knockdown cells for PAK1, PAK2, PPP1R2, and LZTR1, as well as cells ectopically expressing DN PP1C. We then tested their response to the pan-RAS inhibitor RMC-6236^47^. We found that knockdown of PPP1R2 modestly sensitized XG7 and MM.1S cells to RMC-6236, whereas LZTR1 knockdown or expression of DN PP1C conferred resistance (Fig. 6G). Consistent with our previous data, these perturbations had no effect in H1112 cells expressing KRAS A146T or in the PAAD cells. Interestingly, knockdown of either PAK1 or PAK2 sensitized XG7 and MM.1S cells to RAS inhibition, suggesting that both proteins contribute to RAS T148 phosphorylation. Accordingly, we performed an *in vitro* kinase assay with recombinant PAK1 and KRAS. We found that PAK1 could directly phosphorylate KRAS T148, and this activity was suppressed by the PAK1/2 inhibitor FRAX597 (Fig. 6H).

While the PAK1/2 inhibitor FRAX597 alone was not selectively toxic to RAS-dependent MM (Fig. S6I), we hypothesized that it could synergize with a direct RAS inhibitor by lowering oncogenic RAS levels. We combined FRAX597 with RMC-6236 and observed synergistic toxicity in most of the RAS-dependent MM and AML lines tested (Fig. S6K). A notable exception was the H1112 MM line, whose KRAS^A146V^ mutation bypasses the PAK1-LZTR1 degradation axis. We confirmed the mechanism of this drug combination in XG7 cells ectopically expressing KRAS mutants. The combination was effective in cells expressing KRAS^G12D^ but not in cells expressing the KRAS^G12D+T148D^ phosphomimetic that bypasses PAK1 regulation (Fig. 6I).

Finally, we evaluated the PAK1/2 and RAS inhibitor combination in vivo using an MM.1S xenograft model. While single-agent low dose FRAX597 was inactive, RMC-6236 was modestly effective, but the combination significantly suppressed tumor growth without apparent toxicity (Fig. 6J, S6L).

## Discussion

We have uncovered a regulatory circuit governing RAS protein stability, in which phosphorylation of RAS by PAK1 at T148 antagonizes PP1C-mediated dephosphorylation and subsequent LZTR1-dependent degradation (Fig. 6K). This discovery adds a new layer of post-translational control to RAS biology and identifies a previously unknown function of A146 mutations. Our findings establish that regulation of RAS stability by T148 phosphorylation is preferentially active across diverse hematologic malignancies, including MM, AML, and GCB DLBCL, and point to PAK1/2 inhibition as a key therapeutic vulnerability in these cancers.

T148 phosphorylation preferentially protects KRAS and NRAS from degradation in hematologic cancers. This seemingly contrasts with reports from colon cancer models where GSK3B-mediated phosphorylation at T144 and T148 was found to decrease mutant KRAS expression^45,48^. However, we also found that inhibition of GSK3B increased RAS levels (Fig. S6A) and mutation of T144A modestly increased KRAS stability in MM (Fig. 2B-C). We propose that the protective effect of T148 phosphorylation is a dominant regulatory event in MM cells. This difference likely reflects cell-type-specific regulatory mechanisms, potentially linked to distinct upstream signaling networks like the WNT/APC/GSK3B axis dominant in colon cancer^49^. These observations suggest that the unique biology of hematologic malignancies dictates alternative modes of RAS regulation.

The preferential regulation of RAS by PP1C and LZTR1 in hematologic cancers may be caused by multiple factors. One possibility is the differential subcellular localization of RAS signaling. We have previously shown that RAS activates mTORC1 from lysosomes in MM^14^. Since LZTR1 is primarily localized to the Golgi^50^, trafficking between Golgi and endosomes^51^ could expose RAS to LZTR1, thus necessitating T148 phosphorylation to protect RAS. Indeed, LZTR1-associated proteins were enriched for proteins related to vesicle transport (Fig. S3G) and we also observed that PLA puncta between LZTR1 and KRAS^G12D^ were primarily in the cytosol (Fig. 3H). Alternatively, differences in deubiquitinating enzyme (DUB) expression may contribute to this tissue specificity. We found that KRAS was less stable in MM than in colon or pancreatic cells (Fig. 4G, J-K) but that this stability in MM could possibly be increased by overexpression of a DUB not expressed in malignant plasma cells. For instance, the DUB USP18 has been reported to decrease KRAS expression and impede lung tumor growth in mice^52^ but is not expressed in MM patients^38^. Interestingly, this study found that knockdown of USP18 led to a redistribution of KRAS away from the plasma membrane into endomembrane compartments. Perhaps this redistribution allows LZTR1 increased access to KRAS.

Our findings also assign a novel, dual role to KRAS A146 mutations. Beyond their known function in promoting GDP/GTP exchange^39^, we demonstrate that these mutations also protect KRAS from LZTR1-mediated degradation. Given the increased role of LZTR1 as a RAS-regulator in hematologic cancers, this dual function may explain why A146 mutations are enriched in hematologic malignancies but are largely absent from pancreatic and many other epithelial cancers^1^. In support of this notion, experiments in mice found that KRAS^A146T^ induced a myelodysplastic state comparably to KRAS^G12D^ when conditionally expressed in hematopoietic stem cells but failed to effectively drive pancreatic intraepithelial neoplasias (PanINs) or colon crypt hyperplasia^39^. Mechanistically, A146 mutations may obstruct LZTR1 access to the adjacent K147 residue^53^, protecting RAS from ubiquitination and subsequent degradation. Similarly, the negative charge from T148 phosphorylation could also shield K147. Notably, KRAS K147 mutations are not observed in MM^38^, although a recent mutational screen of KRAS noted its oncogenic potential^54^. It is possible that while mutating either A146 or K147 protects KRAS from LZTR1-dependent degradation, only A146 mutations simultaneously enhance GTP cycling, providing potent tandem oncogenic drivers.

Finally, we identify PAK1 and PAK2 as key, previously unrecognized regulators of RAS protein stability. PAK1 and PAK2 are known as downstream effectors of RAC1 and CDC42^55^, and can target MAP kinase, NF-kB, and PI3-K signaling^56^. The role of PAK1 and PAK2 in controlling RAS stability reveals a potential therapeutic vulnerability. We demonstrated that PAK1/2 inhibitors downregulate RAS expression and augment RAS pathway inhibitors, enhancing their efficacy in vivo. We propose that a PAK1-specific inhibitor might improve this synergy by limiting on-target toxicity associated with PAK2 inhibition. Given the limited durability of current KRAS inhibitors, a combination strategy that both inhibits RAS activity and reduces RAS protein levels via PAK1 inhibition may represent a powerful strategy to deepen responses and prevent the emergence of resistance.

## Methods

### Cell culture

MM and adenocarcinoma cell lines were cultured at 37°C in a humidified atmosphere containing 5% CO₂ in Advanced RPMI medium (Invitrogen) supplemented with 4% fetal bovine serum (tetracycline-tested, R&D Systems), 1% penicillin–streptomycin, and 1% L-glutamine (Invitrogen). HEK293FT and HEK293T cells were maintained in DMEM (Invitrogen) supplemented with 10% fetal bovine serum (tetracycline-tested, R&D Systems), 1% penicillin–streptomycin, and 1% L-glutamine (Invitrogen). Cell lines were routinely monitored for mycoplasma contamination using the MycoAlert Mycoplasma Detection Kit (Lonza) and authenticated by short tandem repeat (STR) DNA fingerprinting across 16 copy number variant loci^57^.

| **Name** | **Source** | **Cat. NO.** |
| --- | --- | --- |
| Human: XG-7 | NCI |  |
| Human: RPMI-8226 | ATCC | CCL-155 |
| Human: MM1.S | ATCC | CRL-2974 |
| Human: SK-MM-1 | NCI |  |
| Human: INA-6 | DSMZ | ACC 862 |
| Human: NCI-H929 | ATCC | CRL-3580 |
| Human: THP1 | ATCC | TIB-202 |
| Human: HL-60 | ATCC | CCL-240 |
| Human: AMO1 | DSMZ | ACC 538 |
| Human: H1112 | NCI |  |
| Human: HEK293-FT | Thermo Fisher | R70007 |
| Human: HEK293-T | ATCC | CRL-3216 |
| Human: ASPC1 | ATCC | CRL-1682 |
| Human: PANC-1 | ATCC | CRL-1469 |
| Human: LoVo | ATCC | CCL-229 |
| Human: GP2d | Sigma | 95090714 |

### Antibodies

| **Antibody** | **Species** | **Supplier** | **Catalog No.** |
| --- | --- | --- | --- |
| anti-RAS | rabbit | CST | 91054S |
| anti-PP1C | mouse | Santa Cruz | sc-7482 |
| anti-PPP1R2 | rabbit | R&D | AF4719 |
| anti-PAK1 | rabbit | CST | 2602T |
| anti-PAK2 | rabbit | CST | 2608S |
| anti-LZTR1 | mouse | Santa Cruz | sc-390166 |
| anti-Ubiquitination | mouse | Santa Cruz | sc-8017 |
| anti-SHOC2 | rabbit | CST | 53600S |
| anti-mTOR | mouse | Biolegend | 659202 |
| anti-p-4E-BP1 (S65) | rabbit | CST | 9451S |
| anti-4E-BP1 | rabbit | CST | 9644 |
| anti-MEK1/2 | rabbit | CST | 8727 |
| anti-p-MEK1/2 (S217/221) | rabbit | CST | 9154 |
| anti- GAPDH | mouse | Santa Cruz | sc-47724 |
| anti-rabbit HRP |  | CST | 7074 |
| anti-mouse HRP |  | CST | 7076 |
| anti-goat HRP |  | Santa Cruz | sc-2354 |

### Generation of mNG-Stable Expression Cells

Synthetic KRAS gene fragments (Twist Bioscience) were cloned into the mNG–8xlinker–pBMN–LYT2 vector via Gibson assembly (New England Biolabs) (Table S4). Briefly, 150 ng of KRAS gene fragment was combined with 1 µL of SnaBI-linearized BioID-2 vector and 4 µL of 2× NEBuilder HiFi DNA Assembly Master Mix (New England Biolabs) and incubated for 1 h at 50 °C. The assembled plasmids were transformed into E. coli Stbl3 cells for amplification.

Sequence-verified mNG constructs were packaged into retrovirus using 293T cells (ATCC) together with helper plasmids pHIT60 and pHIT/EA6x3* at a 2:1:1 ratio. Transduced multiple myeloma (MM) cells were sorted on a Sony MA900 cell sorter using an anti-Lyt2 antibody (BioLegend; 1:1000 dilution) in PBS containing 0.5% FBS to establish stable mNG–KRAS-expressing cell lines.

### CRISPR KRAS stability screens

XG7 and RPMI 8226 cells stably expressing mNG-KRAS^G12D^ were transduced in duplicate with the Brunello sgRNA library as described above. Following transduction, cells were selected with puromycin, and Cas9 expression was induced by doxycycline treatment. Cells were expanded for 7 days, after which 5×10⁷ cells were harvested and stored as the baseline (input) sample. The remaining ∼8×10⁷ cells were stained with anti-Lyt2 antibody (BioLegend) at a 1:1000 dilution in PBS supplemented with 0.5% FBS for 20 min on ice. Cells were washed, resuspended at ∼2×10⁷ cells/mL in PBS with 0.5% FBS, and sorted on a Sony MA900 cell sorter to isolate the top and bottom 5% of mNG–KRAS-expressing populations, yielding ∼2×10⁶ cells per fraction. Genomic DNA from sorted populations was extracted using the QIAamp DNA FFPE Advanced Kit (QIAGEN) according to the manufacturer’s protocol.

### CRISPR Library amplification

Genomic DNA from CRISPR screens was processed using a nested PCR strategy. In the first round, sgRNA sequences were amplified from genomic DNA, followed by a second round of PCR to append Illumina-compatible sequencing adapters. Each PCR reaction was performed with ExTaq DNA polymerase (Takara) for 18–20 cycles per round. Amplified products were size-selected using a 2% E-Gel SizeSelect agarose gel (Invitrogen) and quantified with the Qubit dsDNA High-Sensitivity Assay (Thermo Fisher Scientific). Final libraries were sequenced on an Illumina NextSeq 2000 using the NextSeq 1000/2000 Control Software (v1.2.036376).

Demultiplexing was performed with DRAGEN (v3.7.4, Illumina), and sequencing reads were aligned to the reference sgRNA library using Bowtie2 (v2.2.9). PCR primer sequences and detailed protocols are provided in Supplementary Methods.: ^58^.

### CRISPR analysis

The DESeq2 algorithm ^59^ was used to estimate the log-fold change of the read count of sgRNAs between mNG-KRAS high and low samples versus input samples in each experiment. Of the 77,441 guides targeting genes, 9,919 (13%) were removed for having poor performance across a large number of essential gene experiments ^42, 60^. For each gene, the log-ratios of the remaining guides associated with that gene were averaged to estimate a gene-level, log-fold change. For each cell, these gene-level, log-fold changes were normalized by subtracting the mode of their distribution (estimated with the R-function “density”) and then divided by the root-mean- square deviation (RMSD) from that mode. Further analysis was performed using Microsoft Excel v16.63.1.

### Protein interactomes (BioID2)

Synthetic gene fragments of KRAS phospho-mutants, PPP1R2, PP1C, and LZTR1 (Twist) were cloned into the BioID2-10xlinker-pBMN-LYT2 vector ^14^ via Gibson cloning (New England Biolabs) (Table S5). Gene fragments (150 ng) were mixed with 1 μl SnaBI cut BioID-2 vector and 4 μl of the 2x NEBuilder HiFi DNA Assembly Master Mix (New England Biolabs) and incubated for 1 hour at 50°C. The assembled vector was then transformed into Stbl3 cells for amplification. Sequence-verified BioID2 constructs were packaged into retrovirus using 293T cells (ATCC) with helper plasmids pHIT60 and pHIT/EA6x3* in a 2:1:1 ratio. Transduced MM cells were purified with anti-Lyt2 (mouse CD8) magnetic beads (ThermoFisher). Once >95% Lyt2-positive cell pools were obtained, cells were expanded for approximately 2 weeks to obtain 40x10^6^ cells. On the day prior to lysis, biotin (Sigma) was added to a final concentration of 50 μM to BioID2-KRAS, BioID2-PPP1R2, BioID2-PP1C, or LZTR1-BioID2 carrying cells. Cells were incubated for 16 hours in the presence of biotin and then lysed at 1x10^7^ cells per ml in RIPA buffer modified for MS analysis (1% NP-40, 0.5% deoxycholate, 50 mM Tris, pH 7.5, 150 mM NaCl, 1 mM Na_3_VO_4_, 5mM NaF, 1 mM AEBSF) for 10 minutes on ice. Lysates were cleared by centrifugation at 14,000g for 20 minutes at 4°C. Subsequently, 30 μl of pre-washed streptavidin agarose beads (ThermoFisher) was added and samples were rotated at 4°C for 2 hours. At the end of the incubation period, samples were washed two times in 1x RIPA buffer, 1x in PBS, and 1x in 50 mM HEPES and stored at -80°C for later MS analysis. Three independent replicates were collected for all BioID2 experiments.

### Mass spectrometry

For the digestion of KRAS, PPP1R2, PP1C, or LZTR1 BioID2 proteins and TMTpro labeling, each sample bound to capture beads containing 50 µL 50 mM HEPES pH 8 was treated with 150μL of digestion buffer containing the following composition: 10mM TCEP, 40mM chloroacetamide, and 15 ng/µL trypsin/LysC in lysis buffer provided with the EasyPep kit (Thermo Fisher). Samples were incubated at 37°C overnight in the dark at 1000 rpm. After digestion, 175μL of each solution was transferred to a new tube and treated with 10 µL of 10 μg/μL TMTpro (ThermoFisher) reagent and incubated for 1 hour at 25 ⁰C with shaking. Excess TMTpro was quenched with 50μL of 5% hydroxylamine, 20% Formic acid for 10 minutes and samples within each plex were then combined. Combined plexes were cleaned using EasyPep mini columns (ThermoFisher) as described in the manual. Eluted peptides were dried in speed-vac.

For the digestion of global cellular proteins and TMTpro labeling, each cell pellet was lysed in 500 μL EasyPep Lysis buffer (Thermo Fisher) and treated with 1μL universal nuclease (Thermo Fisher). Protein concentration was determined by the BCA method and 10µg was taken from each sample for digestion. Samples were adjusted to 100μL total with lysis buffer and treated with 100μL of digestion buffer containing the following composition: 10mM TCEP, 40mM chloroacetamide, and 15ng/µL trypsin/LysC in 100mM HEPES pH 8. Samples were incubated at 37°C overnight in the dark. Then 10 μL of 10 µg/μL TMTpro 18-plex label (Thermo Fisher) was added to the samples followed by incubation for 1 hour at 25°C. Excess TMTpro was quenched with 50 μL of 5% hydroxylamine, 20% Formic acid for 10 minutes, and samples within each plex were then combined. Samples were cleaned using EasyPep mini columns provided with the EasyPep kit as described in the manual. Eluted peptides were dried in speed-vac.

For LC/MS analysis of BioID2 experiments, peptides were resuspended in 50μL of 0.1% FA and 5μL was analyzed in duplicate using a Dionex U3000 RSLC in front of a Orbitrap Eclipse (Thermo) equipped with an EasySpray ion source. Solvent A consisted of 0.1%FA in water and Solvent B consisted of 0.1%FA in 80%ACN. Loading pump consisted of Solvent A and was operated at 7 μL/min for the first 6 minutes of the run and then dropped to 2 μL/min when the valve was switched to bring the trap column (Acclaim™ PepMap™ 100 C18 HPLC Column, 3μm, 75μm I.D., 2cm) in-line with the analytical column (EasySpray C18 HPLC Column, 2μm, 75μm I.D., 25cm). The gradient pump was operated at a flow rate of 300nL/min and each run used a linear LC gradient of 5-7%B for 1 minute, 7-30%B for 83 minutes, 30-50%B for 25 minutes, 50-95%B for 4 minutes, holding at 95%B for 7 minutes, then re-equilibration of analytical column at 5%B for 17 minutes. All MS injections employed the TopSpeed method with three FAIMS compensation voltages (CVs) and a 1 second cycle time for each CV (3 second cycle time total) that consisted of the following: Spray voltage was 2200V and ion transfer temperature of 300 ⁰C. MS1 scans were acquired in the Orbitrap with resolution of 120,000, AGC of 4e5 ions, and max injection time of 50ms, mass range of 350-1600 m/z; MS2 scans were acquired in the Orbitrap using TurboTMT method with resolution of 15,000, AGC of 1.25e5, max injection time of 22 ms, HCD energy of 38%, isolation width of 0.4Da, intensity threshold of 2.5e4 and charges 2-6 for MS2 selection. Advanced Peak Determination, Monoisotopic Precursor selection (MIPS), and EASY-IC for internal calibration were enabled and dynamic exclusion was set to a count of 1 for 15 seconds.

For LC/MS analysis of cellular proteomes, global cellular peptides were resuspended in 50μL of 0.1% FA and loaded 5µL twice onto a Dionex U3000 RSLC in front of a Orbitrap Eclipse (Thermo) equipped with an EasySpray ion source. Solvent composition and column configuration was the same as described in the previous section. The gradient pump was operated at a flow rate of 300nL/min and each run used a linear LC gradient of 5-7%B for 1min, 7-30%B for 133 minutes n, 30-50%B for 35 minutes, 50-95%B for 4 minutes, holding at 95%B for 7 minutes, then re-equilibration of analytical column at 5%B for 17 minutes. All MS injections employed the TopSpeed method with four FAIMS compensation voltages (CVs) and a 0.75 second cycle time for each CV (3 second cycle time total) that consisted of the following: Spray voltage was 2200V and ion transfer temperature of 300 ⁰C. MS1 scans were acquired in the Orbitrap with resolution of 120,000, AGC of 4e5 ions, and max injection time of 50ms, mass range of 350-1600 m/z; MS2 scans were acquired in the Orbitrap using TurboTMT method with resolution of 15,000, AGC of 1.25e5, max injection time of 22ms, HCD energy of 38%, isolation width of 0.4Da, intensity threshold of 2.5e4 and charges 2-6 for MS2 selection. Advanced Peak Determination, Monoisotopic Precursor selection (MIPS), and EASY-IC for internal calibration were enabled and dynamic exclusion was set to a count of 1 for 15 seconds.

### shRNA and sgRNA mediated knockdown

Individual shRNAs were obtained from the MISSION shRNA Library from the RNAi Consortium TRC1.0 in the pLKO.1 vector (SIGMA):

shCTRL GCCAAGATTCAGAATCCCAAA

shPPP1R2.1 CTACATGATGATGATGAAGAT

shPPP1R2.2 CCAGACATCTTAGCCAGGAA

shPPP1CA.1 CCGCAATTCCGCCAAAGCCAA

shPPP1CA.2 GCTGCTGGCCTATAAGATCAA

shSHOC2.1 CAATACGATCAAACGGCCAAA

shSHOC2.2 CATGCTTAGCATTCGAGAGAA

shLZTR1.1 GCACATCATTGTGCACCAGTTCTC

shLZTR1.2 CTTCAAGAAGTCCCGAGATGTCTC

Lentiviral packaging and transduction of shRNAs and sgRNAs were performed as described above. Briefly, shRNA and sgRNA vectors were packaged in 293FT cells (Invitrogen) using the helper plasmids pPAX2 (Addgene 12260) and pMD2.g (Addgene 12259) at a 4:3:1 ratio.

Supernatants from the 293FT cells were collected at 24 and/or 48 hours post-transfection. MM cells were transduced for 90 minutes at 2500 rpm at 30°C, supplemented with fresh complete media, and incubated overnight before subsequent experiments. For shRNA experiments, transduced MM cells were subjected to selection with 1 μg/ml puromycin (Gibco) for 2 days, followed by Western blot analysis. Individual shRNAs were acquired from the MISSION shRNA Library, developed by the RNAi Consortium, and cloned into the pLKO.1 vector (Sigma). For sgRNA experiments, transduced MM cells were either left untreated or selected using 1 μg/ml puromycin. sgRNAs were cloned into the pLKO.1-puro-U6 or pLKO.1-puro-GFP vector for subsequent analysis.

### Site-directed mutagenesis

Set up PCR reaction using PrimeSTAR HS DNA polymerase (TaKaRa) to amplify the original plasmid with the designed primers. Following with cycling conditions: initial denaturation: 98°C, 3 minutes; 35 cycles of denaturation 98°C 10 seconds, annealing 55°C 5 seconds, extension 72°C 8 minutes; final extension 72°C 10 minutes. Then, purify the PCR product using DNA Clean&Concentrator (ZYMO Research). Purified product were digested using DpnI (NEB). Transform the DpnI-treated DNA into competent E.coli cells. All mutations were verified by whole plasmid Nanopore sequencing (Quintarabio). All KRAS mutant sequences are found in Table S5.

### Western blot analysis

Cells were lysed at a 1x10^7^ cells per mL in modified RIPA buffer (1% NP-40, 0.5% deoxycholate, 50 mM Tris, pH 7.5, 150 mM NaCl, 1 mM Na3VO4, 5 mM NaF, 1 mM AEBSF) for 30 minutes on ice. Lysates were cleared by centrifugation at 14,000g for 20 minutes at 4 °C, and the post-nuclear supernatant was collected. Protein concentrations were determined using the Pierce BCA protein assay kit (Thermo) following the manufacturer’s protocol. A total of 100 µL of lysate was mixed with 40 µL of 4x Laemmli sample buffer (Bio-Rad) containing 1% β-mercaptoethanol (Bio-Rad) and boiled for 10 minutes. Subsequently, 15 µg of each lysate was loaded onto a 4–12% gradient gel (Bio-Rad) and transferred to a PVDF membrane (Millipore) using a Bio-Rad Transblot Turbo semi-dry transfer device (Bio-Rad). PVDF membranes were blocked with 5% milk (Bio-Rad) in TBST and subsequently probed with the indicated antibodies. Phospho-specific antibodies were diluted in 1% BSA (MPI), while other antibodies were diluted in milk. Detection was performed using HRP-conjugated secondary antibodies (anti-rabbit-HRP or anti-mouse-HRP; Cell Signaling Technology) as appropriate. Chemiluminescent signals were visualized using the ChemiDoc Imaging System (Bio-Rad) and analyzed with Image Lab Touch Software (v2.3.0.07) (Bio-Rad).

### Proximity Ligation Assay

MM cells were plated onto a 15 well m-Slide Angiogenesis ibiTreat chamber slide (Ibidi) and allowed to adhere to the surface for 30 minutes at 37°C. Cells were next fixed with 4% paraformaldehyde (Electron Microscopy Sciences) for 20 minutes at room temperature and then washed in PBS (Gibco). Cells were permeabilized in cold methanol for 20 minutes at -20°C, washed in PBS and then blocked in Duolink Blocking buffer (Sigma) for 30 minutes at room temperature. Primary antibodies were diluted in Duolink Antibody Diluent (Sigma) and incubated overnight at 4°C. Cells were next washed for 2x 4 minutes in TBST with 0.05% tween-20, followed by addition of the appropriate Duolink secondary antibodies (Sigma), diluted and mixed according to the manufacturer’s instructions. Cellular membranes were labeled by the addition of 5 mg/ml wheat germ agglutinin (WGA) conjugated to Alexa Fluor 488 (Thermo Fisher), and cells were incubated for 1 hour at 37°C, after which the plates were washed in TBST with 0.05% tween-20 for 2x5 minutes. Ligation and amplification steps of the PLA were performed using the Duolink in situ Detection Reagents Orange kit (Sigma) according to the manufacturer’s instructions. Following the PLA, cells were mounted in Fluoroshield Mounting Medium with DAPI (Abcam). Images were acquired on a Leica Stellaris 5 Confocal microscope using Leica Application Suite X version 4.5.0.2531. Images for display were prepared with NIH ImageJ/FIJI software version 2.9.0/1.53t ^61^.

### PLA image analysis

PLA puncta were counted using custom macro available at https://github.com/janwisn/EIB_PLA_Analysis. PLA Score was determined by normalizing the number of PLA spots counted in each sample to the average number of PLA spots counted in the control sample, which was set to 100. Box and whisker plots display the median PLA Score with whiskers incorporating 10-90% of all data; outliers are displayed as dots. Statistical comparisons were made by one-way ANOVA with Dunnett’s post test using Prism.

### FACS analysis

For FACS analysis, mutant RAS isoforms were linked to mNeonGreen as gene fragments (Twist Biosciences) and stained with anti-CD8 (1:2000) as previously described (PMID: 29925955). mutant RAS expression in MM was measured by staining 2x10^5^ cells on ice with anti-CD8 (1:2000) for 20 minutes in FACS buffer (PBS with 2% BSA). Cells were washed with FACS buffer for twice and resuspended in 250μl of FACS buffer. These cells were analyzed on a Beckman Coulter Cytoflex using CytExpert software and analyzed with FlowJo version 10.

### Co-immunoprecipitation

For co-immunoprecipitation, mutant RAS isoforms were linked to mNeonGreen as gene fragments (Twist Biosciences). MM cells were retrovirally transduced and selected for mNeonGreen expression. For lysis, 20x10^6^ cells were lysed in either 0.5% CHAPS for co-immunoprecipitation (co-IP) of RAS in lysis buffer composed of 50 mM Tris (pH 7.5), 150 mM NaCl, 1 mM Na3VO_4_, 5 mM NaF, and 1 mM AEBSF. Lysis was carried out on ice for 10 minutes, followed by clearing the lysates through centrifugation at 14,000g for 20 minutes at 4°C. The supernatant was collected. The lysates were divided into two portions, and 25 μL of mNeonGreen-Trap agarose (Chromotek) was added to pull down mNeonGreen-tagged RAS constructs, while a control sample received 25 μL of saturated control beads (Chromotek).

Lysates were rotated at 4°C for 2 hours to allow binding. Subsequently, beads were washed three times with CHAPS lysis buffer. To each sample, 30 μL of 2x Laemmli sample buffer (BioRad) was added, followed by a 5 minutes incubation at 95°C. The samples were then subjected to western blot analysis as outlined above.

### Immunoprecipitation

Same as co-immunoprecipitation, in immunoprecipitation mutant RAS isoforms were linked to mNeonGreen as gene fragments (Twist Biosciences). 20 x10^6^ selected MM cells were lysed in 1% SDS buffer in PBS for 10 minutes, followed by 30s sonicate/30s break. Then, the samples were boiled at 95°C for 10 minutes. Dilute the lysates from 1% SDS with 1% NP40 to 0.1% SDS. The samples were then subjected to mNeonGreen-Trap agarose purification and western blot analysis as described below.

### Pathway analysis

Pathway analysis was performed using ToppFun from the ToppGene Suite (https://toppgene.cchmc.org)^62^. Gene Ontology^34^ Biological Process analysis was used with the LZTR1-BioID2 dataset as indicated in text.

### Alphafold3 protein-protein interaction prediction

The protein structure complexes were predicted using AlphaFold3 ^37^. Full-length LZTR1 (UniProt: Q8N653), KRAS (UniProt: P01116), CUL3 (UniProt: Q13618) and the ligand GDP (Guanosine-5’-diphosphate) were used as inputs for complex prediction. The variants KRAS^G12D^, KRAS^G12D+T148A^, and KRAS^G12D+T148D^ represent full-length KRAS proteins carrying the G12D with additional substitutions at residue 148 as indicated. The interface predicted TM-score (ipTM) was obtained directly from AlphaFold3 predictions, while the predicted aligned error (PAE) was visualized using the PAE Viewer ^63^. The protein complex structures were visualized and analyzed using ChimeraX software ^64^.

### Viability assays

MM cell lines were seeded at densities of 2,000 to 10,000 cells per well in triplicate in 96-well plates, with three independent biological replicates for each cell line. FRAX597, RMC-6236 (SelleckChem) were dissolved in DMSO and diluted in equal volumes at the specified concentrations. Cells were treated with the compounds for 3 to 7 days, with drug replenishment occurring after 3 days. Metabolic activity was assessed at the end of the incubation period using the CellCountingKit-8 (SelleckChem) according to the manufacturer’s instructions. Absorbance was measured at 450 nm using a Tecan Infinite 200 Pro plate reader.

### Xenograft experiments

All mouse experiments were approved by the National Cancer Institute Animal Care and Use Committee (NCI-ACUC) and were performed in accordance with NCI-ACUC guidelines and under approved protocols. Female NSG (non-obese diabetic/severe combined immunodeficient/common gamma chain deficient) mice were obtained from NCI Fredrick Biological Testing Branch and used for the xenograft experiments between 6-8 weeks of age. Mice were housed in specific pathogen-free facility in ventilated microisolator cages with 12 hour light and 12 hour dark cycles at 72F and 40-60% relative humidity. Approved protocols allowed tumor growth below 20 mm in any dimension; no animals had tumors which exceeded these limits. MM.1S multiple myeloma tumors were established by subcutaneous injection of 10^6^ cells in a PBS. Treatments were initiated when tumor volume reached a mean of 100 mm^3^.

RMC-6236 and FRAX597 (MedChemExpress) were prepared in sterile-filtered 20% (2-Hydroxypropyl)-β-cyclodextrin (Sigma) in PBS and administered p.o. once per day (5mg/kg/day for RMC-6236, 3mg/kg/day or 3 mg/kg/day for FRAX597). For the combination arm, drugs were given at the same concentration and schedule as single agents. Each treatment group contained 5 mice. Tumor growth was monitored every other day by measuring tumor size in two orthogonal dimensions, and tumor volume was calculated by the following equation: tumor volume = (length × width^2)/2.

## Data availability

Analyzed data from all CRISPR screens are provided in Supplemental Table 1 and sequencing reads for all CRISPR screens are made available through our website at https://lymphochip.nih.gov/local/RAS_stability. This paper also analyzes existing, publicly available data: CHRONOS scores were obtained from the DepMap portal (https://depmap.org/portal/) using the 23Q4 data release. Mass spectrometry data was uploaded to Mass Spectrometry Interactive Virtual Environment (MassIVE) under accession number MSV000097703. Analyzed proteomics datasets are available in Supplemental Tables 2-3.

Public MM patient RNA-seq and whole exome sequencing data were obtained from the MMRF-CoMMpass study (IA17 release) and are available at https://research.themmrf.org.

## Acknowledgements

This research was supported by the Intramural Research Program of the NIH, Center for Cancer Research, National Cancer Institute (R.M.Y.). We thank Drs. Lou Staudt, Dan Hodson, Jagan Muppidi, Constantine Mitsiades, and Ji Luo for informative discussions and Dr. Xin-Shuan Su for reagents.

## Author Contributions

Conceptualization: L.Z., R.M.Y.; Experimental design: L.Z., A.B., O.O., C.V.W.; Proteomics: R.J.H., T.A.; Wrote the manuscript: L.Z., R.M.Y.

The authors declare no potential conflicts of interest.

