## Supplemental Figures for "Phosphorylation Protects Oncogenic RAS from LZTR1-Mediated Degradation"

**Supplementary Figures and Legends**

### Figure S1

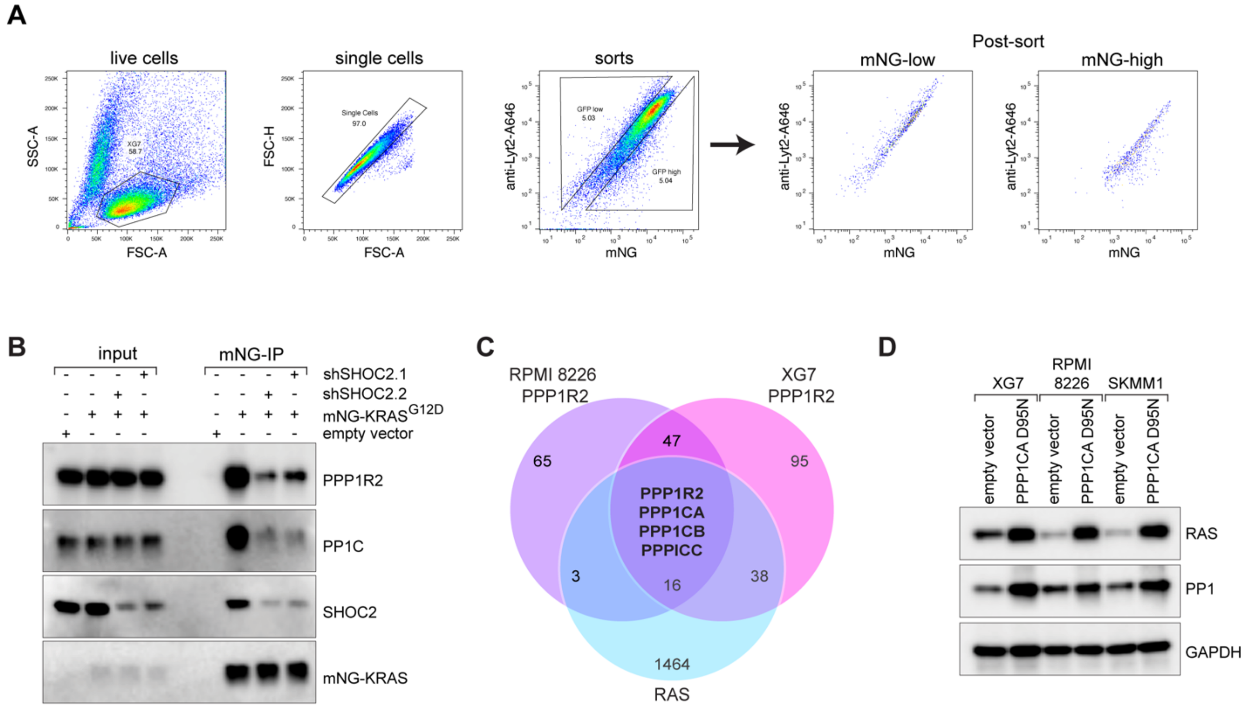


**Supplementary Figure 1. PPP1R2 and PP1C regulate RAS stability. A)** FACS sorting workflow for mNG-KRAS-low and -high cells. **B)** Co-IP with western blot analysis of $KRAS^{G12D}$ pulldown with PPP1R2, PPP1C and SHOC2 following transduction with shCTRL or two SHOC2 shRNAs, n=2. **C)** Venn diagram of overlapping genes from RAS BioID2 proteomics (≥1.0 log2 enrichment vs. control (*5*)), PPP1R2 BioID2 proteomics (≥1.0 log2 enrichment vs. control) in XG7 and RPMI 8226 cells. **D)** Immunoblot analysis of RAS, PP1C and GAPDH after transduction with empty vector or DN PP1C in XG7, RPMI 8226 and SKMM1 cells, n=4.

### Figure S2

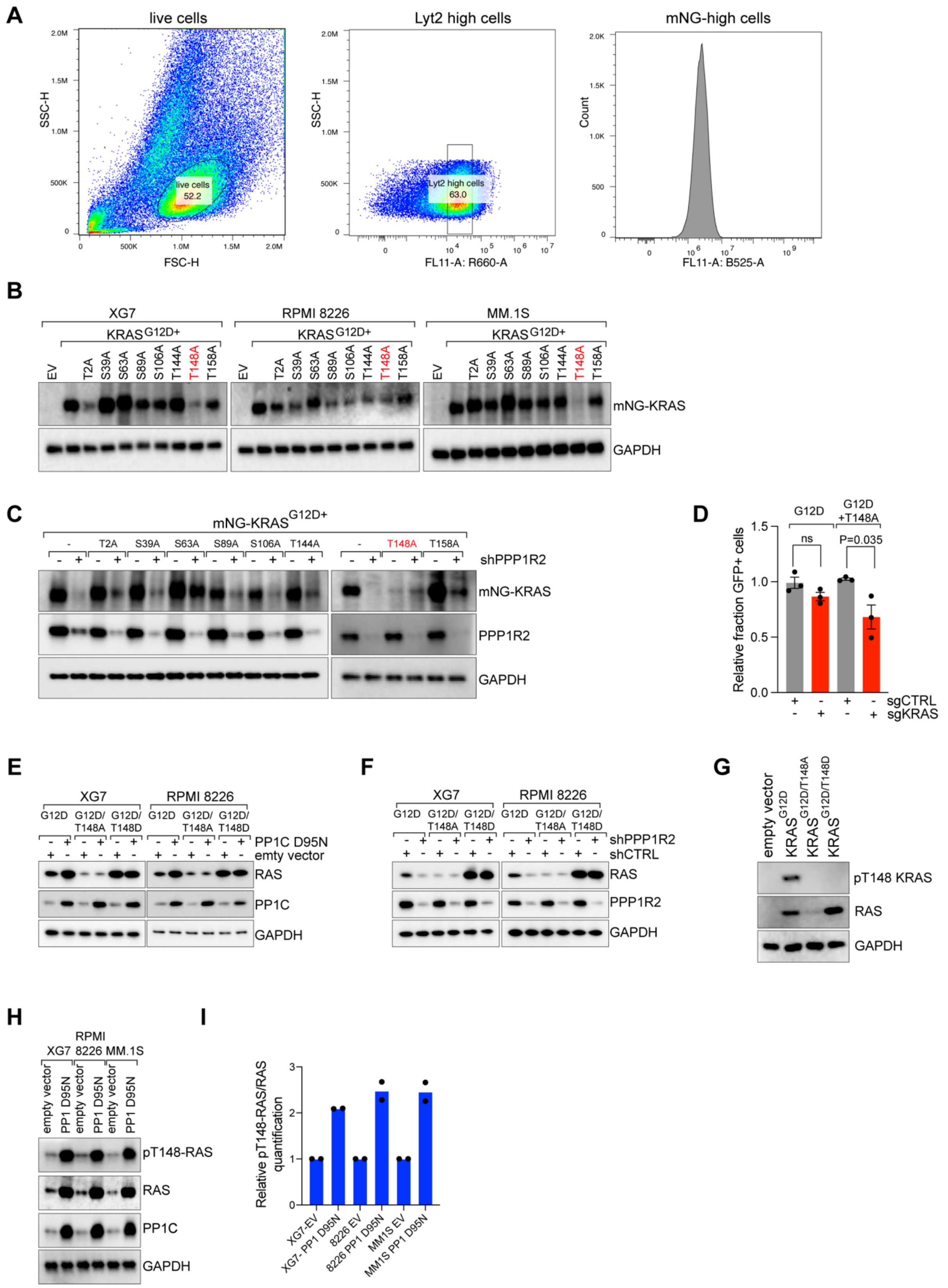
A
live cells
Lyt2 high cells
mNG-high cells
SSC-H
FSC-H
FL11-A: R660-A
Count
FL11-A: B525-A
live cells 52.2
Lyt2 high cells 63.0
B
XG7
RPMI 8226
MM.1S
KRAS G12D+
EV
T2A
S39A
S63A
S89A
S106A
T144A
T148A
T158A
mNG-KRAS
GAPDH
C
mNG-KRAS G12D+
shPPP1R2
PPP1R2
D
G12D
G12D +T148A
ns
P=0.035
Relative fraction GFP+ cells
sgCTRL
sgKRAS
E
G12D/ T148A
G12D/ T148D
PP1C D95N
emty vector
RAS
PP1C
F
shCTRL
G
empty vector
KRAS G12D
KRAS G12D/T148A
KRAS G12D/T148D
pT148 KRAS
H
XG7
RPMI 8226
MM.1S
PP1 D95N
pT148-RAS
I
Relative pT148-RAS/RAS quantification
XG7-EV
XG7- PP1 D95N
8226 EV
8226 PP1 D95N
MM1S EV
MM1S PP1 D95N

**Supplementary Figure 2. T148 of KRAS is directly targeted by PP1C. A)** FACS analysis workflow to determine mNG-KRAS expression of various phospho-mutants. **B)** Immunoblot analysis of mNG-$KRAS^{G12D}$ and GAPDH in XG7, RPMI 8226, and MM.1S cells harboring indicated mutations, n=3. **C)** Western blot analysis for mNG-KRAS, PPP1R2, and GAPDH in XG7 cells expressing indicated mNG-KRAS mutants and either shCTRL or shPPP1R2.1. **D)** Average normalized $KRAS^{G12D}$ or $KRAS^{G12D+T148A}$ CRISPR-mediated rescue viability data following 12 days of KRAS knockout, n=3, error bars depict SEM. **E)** Western blot analysis of RAS, PP1C and GAPDH with empty vector or DN PP1C in XG7 and RPMI 8226 cells expressing $KRAS^{G12D}$, $KRAS^{G12D+T148A}$,or $KRAS^{G12D+T148D}$, n=2. **F)** Western blot analysis of RAS, PPP1R2 and GAPDH transduced with shCTRL or shPPP1R2.1 in XG7 and RPMI 8226 cells expressing $KRAS^{G12D}$, $KRAS^{G12D+T148A}$,or $KRAS^{G12D+T148D}$, n=2. **G)** Western blot analysis of pT148 KRAS, RAS and GAPDH in XG7 cells expressing empty vector, $KRAS^{G12D}$, $KRAS^{G12D+T148A}$,or $KRAS^{G12D+T148D}$, n=2. **H)** Western blot analysis of pT148 KRAS, RAS, PP1C and GAPDH in XG7, RPMI 8226 and MM.1S cells expressing empty vector, or DN PP1C, n=2 **I)** Ratio of pT148-RAS to RAS from quantified blots (from Figure S2H) in XG7, RPMI 8226, and MM.1S cells expressing empty vector or DN PP1C.

### Figure S3

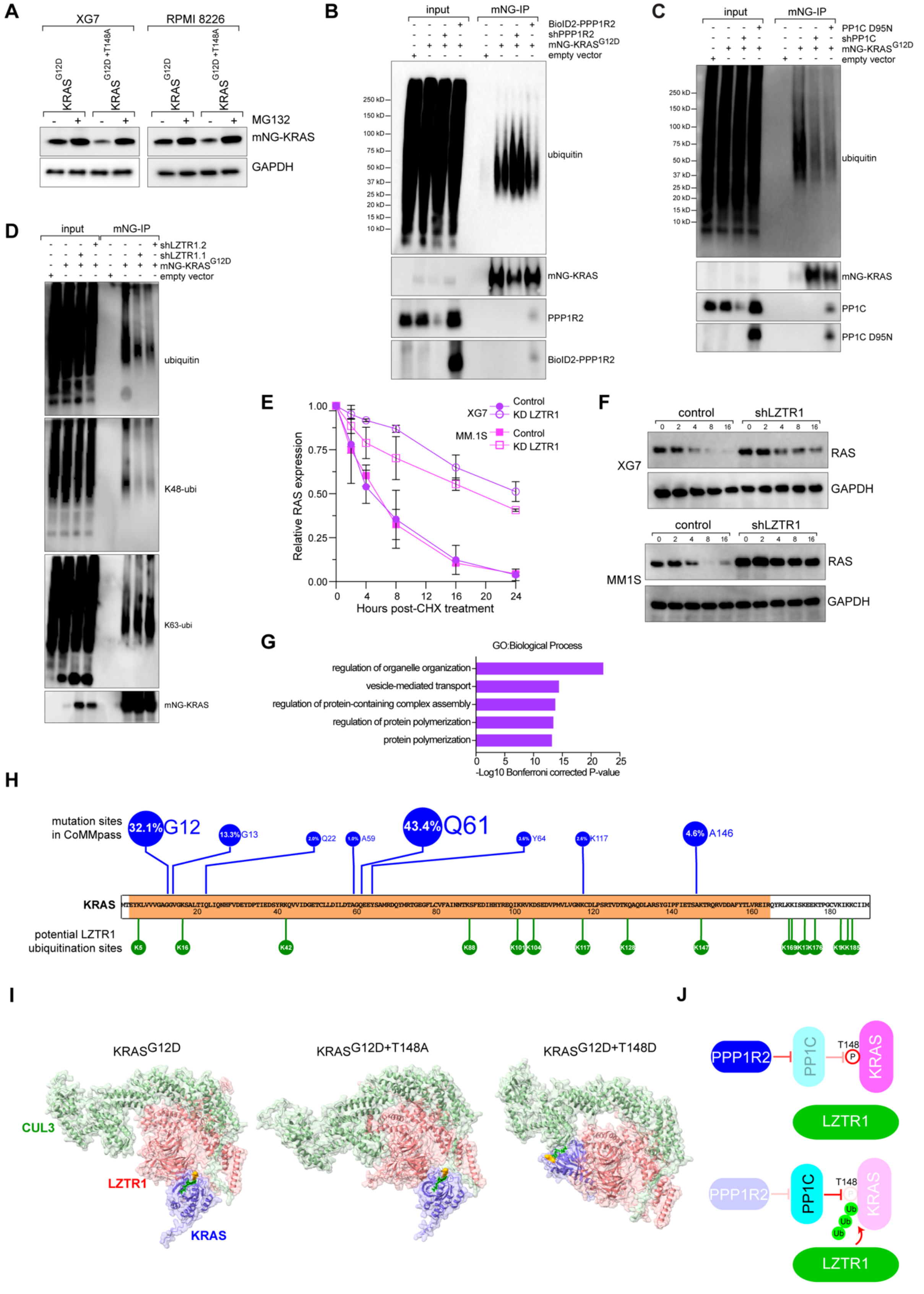
A
XG7
RPMI 8226
MG132
mNG-KRAS
GAPDH
B
input
mNG-IP
BioID2-PPP1R2
shPPP1R2
mNG-KRAS G12D
empty vector
ubiquitin
mNG-KRAS
PPP1R2
BioID2-PPP1R2
C
input
mNG-IP
PP1C D95N
shPP1C
mNG-KRAS G12D
empty vector
ubiquitin
mNG-KRAS
PP1C
PP1C D95N
D
input
mNG-IP
shLZTR1.2
shLZTR1.1
mNG-KRAS G12D
empty vector
ubiquitin
K48-ubi
K63-ubi
mNG-KRAS
E
Relative RAS expression
Hours post-CHX treatment
XG7
Control
KD LZTR1
MM.1S
Control
KD LZTR1
F
control
shLZTR1
XG7
RAS
GAPDH
MM1S
RAS
GAPDH
G
GO:Biological Process
regulation of organelle organization
vesicle-mediated transport
regulation of protein-containing complex assembly
regulation of protein polymerization
protein polymerization
-Log10 Bonferroni corrected P-value
H
mutation sites in CoMMpass
32.1% G12
13.3% G13
Q22
A59
43.4% Q61
Y64
K117
4.6% A146
KRAS
potential LZTR1 ubiquitination sites
I
KRAS G12D
KRAS G12D+T148A
KRAS G12D+T148D
CUL3
LZTR1
KRAS
J
PPP1R2
PP1C
T148
KRAS
LZTR1
Ub

**Supplementary Figure 3. PPP1R2 and PP1C regulate RAS ubiquitination modification. A)** Western blot analysis of mNG-KRAS$^{G12D}$, mNG-KRAS$^{G12D+T148A}$,and GAPDH in XG7 and RPMI 8226 cells treated with or without 10 nM MG132 for 8 hours, n=3. **B)** Western blot analysis of ubiquitin binding following mNG-KRAS pulldown in cells transduced with empty vector, shPPP1R2.1, or BioID2-PPP1R2, n=2. **C)** Western blot analysis of ubiquitin binding following mNG-KRAS pulldown in cells transduced with empty vector, shPP1C, or DN PP1C, n=3. **D)** Western blot analysis of total, K48, and K63 ubiquitin binding following mNG-KRAS pulldown in cells transduced with empty vector or shLZTR1, n=2. **E)** Quantification of immunoblots of RAS expression normalized to GAPDH from XG7 and MM.1S with shCTRL or shLZTR1 following a time course of treatment with 10 nM cycloheximide (CHX) for the indicated timepoints (n=2; error bars depict standard deviation; representative blots in Fig. S3F). **F)** Representative western blots from panel E. G) Bar graph of Gene Ontology enrichment from LZTR1-BioID2 experiment. **H)** Schematic of KRAS oncogenic hotspot mutations and putative lysine ubiquitination sites. **I)** AlphaFold modeling of KRAS$^{G12D}$ with additional T148A or T148D mutation to model a negative charge at this position. K147 is highlighted in yellow and GDP is in green. **J)** Model of T148-dependent regulation of RAS protein stability, in which phosphorylation at T148 protects KRAS from LZTR1-mediated degradation.

### Figure S4

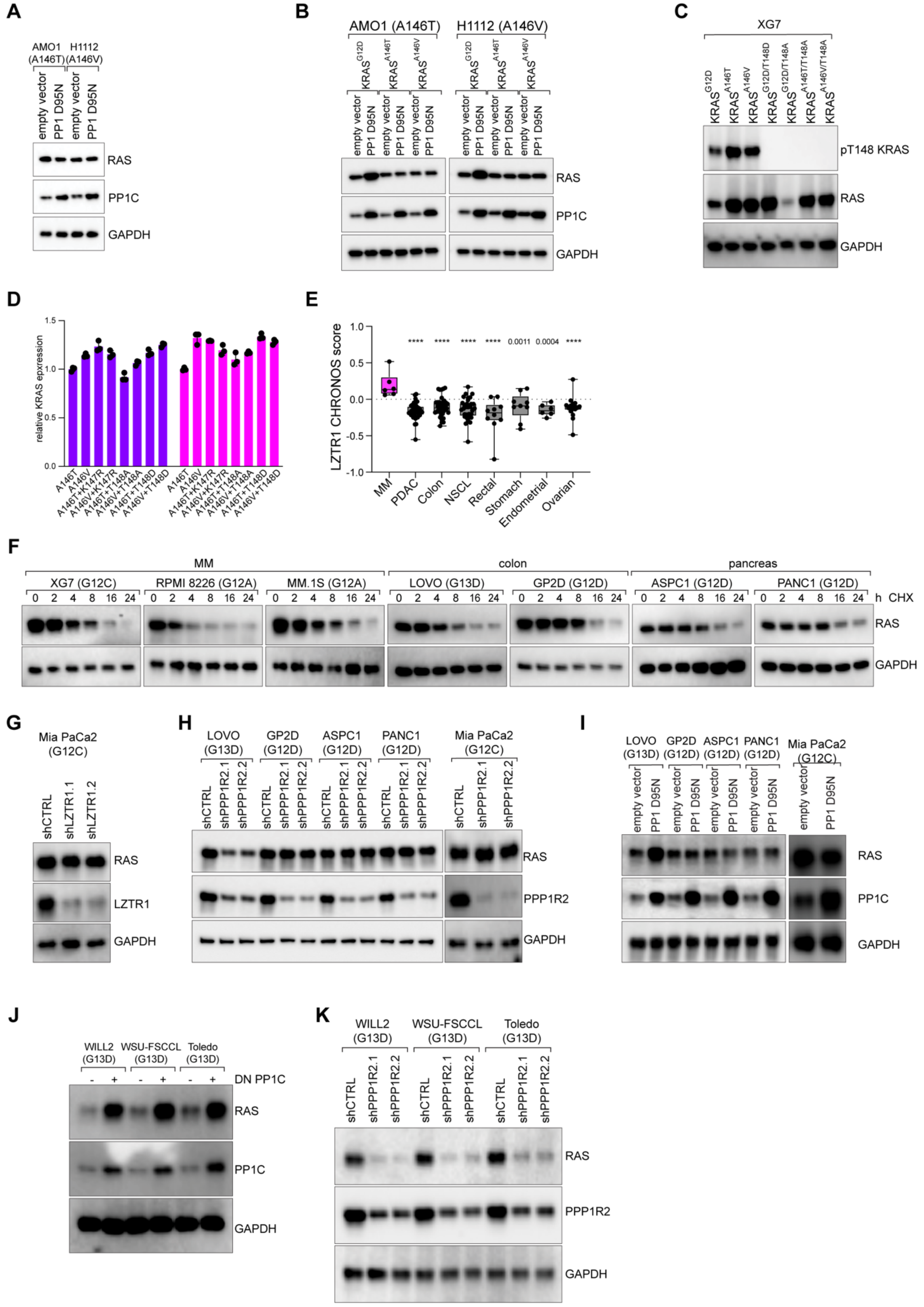

A
AMO1 (A146T)
H1112 (A146V)
empty vector
PP1 D95N
RAS
PP1C
GAPDH
B
AMO1 (A146T)
H1112 (A146V)
KRASG12D
KRASA146T
KRASA146V
empty vector
PP1 D95N
RAS
PP1C
GAPDH
C
XG7
KRASG12D
KRASA146T
KRASA146V
KRASG12D/T148D
KRASG12D/T148A
KRASA146T/T148A
KRASA146V/T148A
pT148 KRAS
RAS
GAPDH
D
relative KRAS epxression
A146T
A146V
A146T+K147R
A146V+K147R
A146T+T148A
A146V+T148A
A146T+T148D
A146V+T148D
E
LZTR1 CHRONOS score
****
0.0011
0.0004
MM
PDAC
Colon
NSCL
Rectal
Stomach
Endometrial
Ovarian
F
MM
colon
pancreas
XG7 (G12C)
RPMI 8226 (G12A)
MM.1S (G12A)
LOVO (G13D)
GP2D (G12D)
ASPC1 (G12D)
PANC1 (G12D)
h CHX
RAS
GAPDH
G
Mia PaCa2 (G12C)
shCTRL
shLZTR1.1
shLZTR1.2
RAS
LZTR1
GAPDH
H
LOVO (G13D)
GP2D (G12D)
ASPC1 (G12D)
PANC1 (G12D)
Mia PaCa2 (G12C)
shCTRL
shPPP1R2.1
shPPP1R2.2
RAS
PPP1R2
GAPDH
I
LOVO (G13D)
GP2D (G12D)
ASPC1 (G12D)
PANC1 (G12D)
Mia PaCa2 (G12C)
empty vector
PP1 D95N
RAS
PP1C
GAPDH
J
WILL2 (G13D)
WSU-FSCCL (G13D)
Toledo (G13D)
DN PP1C
RAS
PP1C
GAPDH
K
WILL2 (G13D)
WSU-FSCCL (G13D)
Toledo (G13D)
shCTRL
shPPP1R2.1
shPPP1R2.2
RAS
PPP1R2
GAPDH

**Supplementary Figure 4. A146 mutations protect KRAS from LZTR1. A)** Western blot analysis of RAS, PP1C, and GAPDH in AMO1 and H1112 lines following expression of empty vector or DN PP1C, n=3. **B)** Western blot analysis of RAS, PP1C, and GAPDH after transducing empty vector or DN PP1C in AMO1 and H1112 cells expressing empty vector, $KRAS^{G12D}$, $KRAS^{A146T}$,or $KRAS^{A146V}$, n=2. **C)** Western blot analysis of pT148 KRAS, RAS and GAPDH in XG7 cells expressing empty vector, $KRAS^{G12D}$, $KRAS^{A146T}$, $KRAS^{A146V}$, $KRAS^{A146T+T148A}$,or $KRAS^{A146V+T148A}$, n=2. **D)** Average normalized mNG-KRAS mutations expressing shCTRL or shLZTR.1 from FACS analysis, n=3, error bars depict standard deviation. **E)** Box plot of KRAS LZTR1 CHRONOS score from depmap for indicated tumor types, with MM highlighted in pink. **F)** Immunoblot analysis of RAS expression within indicated cell lines following treatment with 10 nM cycloheximide (CHX) for the indicated timepoints, n=2. **G)** Immunoblots of RAS, LZTR1, and GAPDH following expression of control or LZTR1 shRNAs in Mia PaCa2, n=2. **H-I)** Western blot analysis of RAS expression following transduction with control shRNA PPP1R2 (**H**) or following ectopic expression of DN PP1C **(I)**, n=3. **J-K**) Western blot analysis of RAS expression in WILL2, WSU-FSCCL, or Toledo GCB DLBCL cell lines following ectopic expression of DN PP1C (**J**), or following transduction with control shRNA or shRNAs targeting PPP1R2 (**K**), (n=3).

### Figure S5

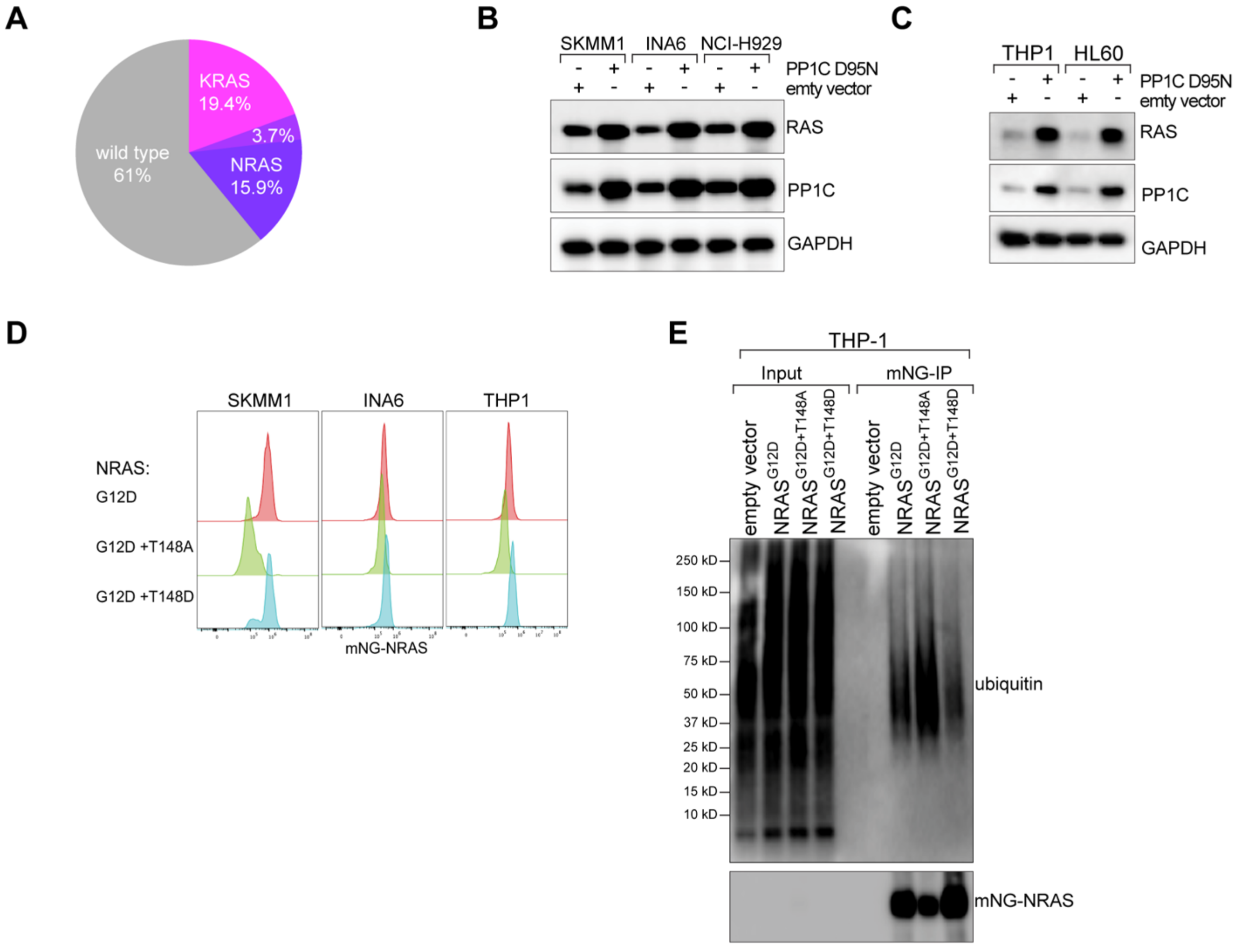


**Supplementary Figure 5. PPP1R2 and PP1C control NRAS stability. A)** Pie chart showing RAS mutation distribution in MM from MMRF CoMMpass data for tumors harboring KRAS (pink), NRAS (purple), both KRAS and NRAS (light purple), or wild-type RAS (gray). **B**) Western blot analysis of RAS, PP1C, and GAPDH with empty vector or DN PP1C in SKMM1, INA6, and NCI-H929 cells (n=3). **C**) Western blot analysis of RAS, PP1C, and GAPDH with empty vector or DN PP1C in THP1 and HL60 cells (n=3). **D**) Representative FACS data of mNG-KRAS$^{G12D}$, KRAS$^{G12D+T148A}$,or KRAS$^{G12D+T148D}$, n=2. **E)** Western blot analysis of ubiquitin binding following mNG-NRAS pulldown in cells transduced with indicated NRAS mutants in THP1 AML cells, n=3.

### Figure S6

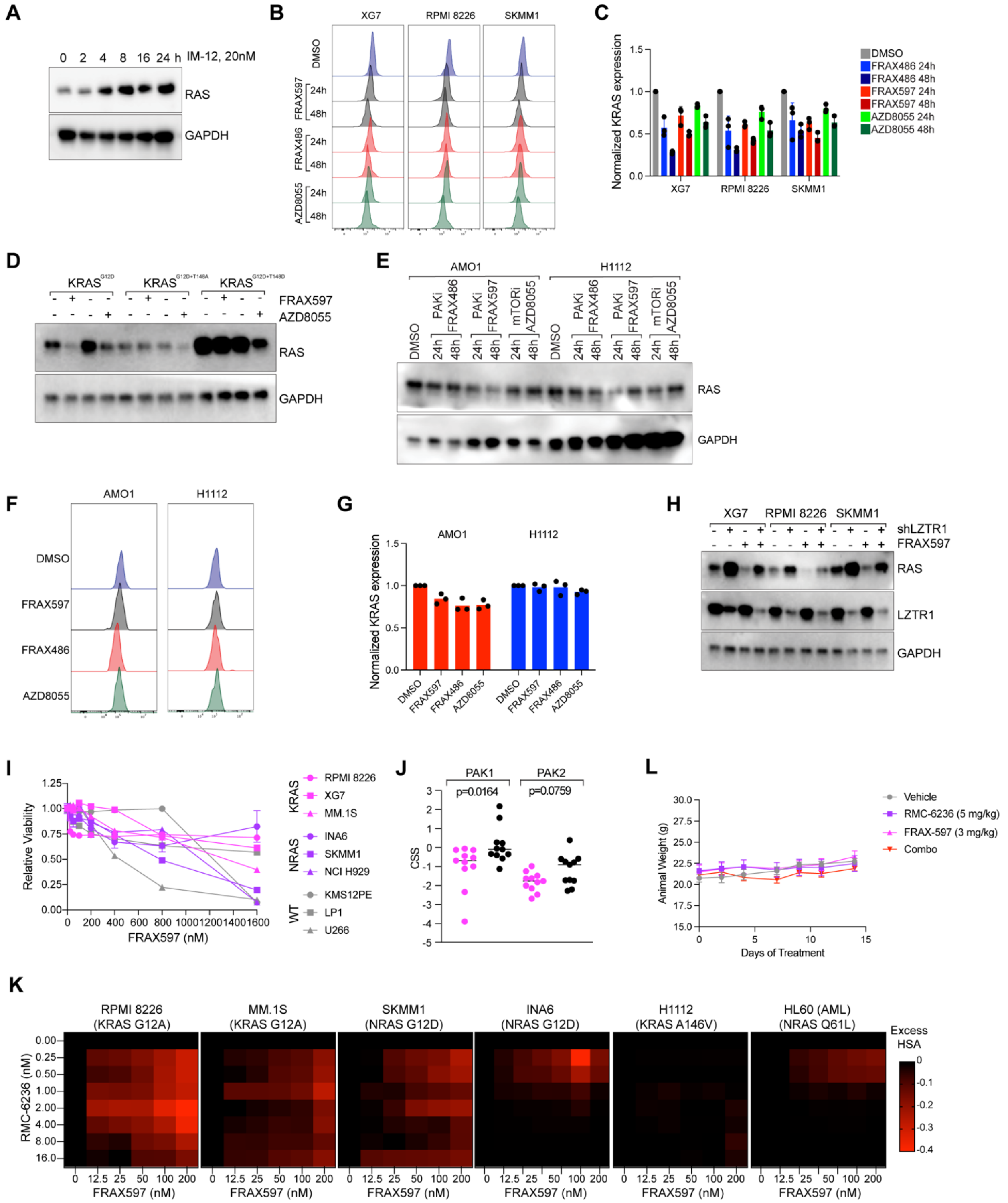


**Supplementary Figure 6. PAK1/2 inhibition blocks T148 phosphorylation. A)** Western blot analysis of RAS and GAPDH after treatment with GSK3B inhibitor, IM-12, at 20 nM for 2, 4, 8, 16, and 24 hours (n=2). **B**) Representative FACS data of mNG-KRASG12D after treatment with

PAKi (FRAX597 500 nM, FRAX486 500 nM, AZD8055 100 nM) for 24 and 48 hours (n=3).**C**) Quantification of FACS data from (B) normalized to DMSO for indicated drugs and cell lines (n=3; error bars depict standard deviation). **D)** Western blot analysis of RAS and GAPDH in XG7 cells expressing $KRAS^{G12D}$, $KRAS^{G12D+T148A}$,or $KRAS^{G12D+T148D}$ after treated with 500 nM FRAX597 or 100 nM AZD8055 for 24 h. **E)** Western blot analysis of RAS and GAPDH following treatment with FRAX597 500 nM, FRAX486 500 nM, and AZD8055 100 nM for 24 and 48 hours in AMO1 and H1112 cells. **F**) Representative FACS data of mNG-KRASG12D after treatment with PAKi (FRAX597 500 nM, FRAX486 500 nM, AZD8055 100 nM) for 48 hours in AMO1 and H1112 cells. **G**) Average normalized FACS data from mNG-KRASG12D with indicated inhibitor treatment from (E) in AMO1 and H1112 cells (n=3; error bars depict standard deviation). **H)** Western blot analysis of RAS, LZTR1, and GAPDH in XG7, RPMI 8226, and SKMM1 cells treated with 500 nM FRAX597 for 24 hours and expressing shCTRL or shLZTR1 (n=4).**I**) Cell viability following 4 days of treatment with FRAX597 at the indicated doses and in the indicated KRAS-dependent, NRAS-dependent, and RAS-independent MM lines, n=2. **J**) CRISPR Screen Scores (CSS) for PAK1 and PAK2 in RAS-dependent (pink) or RAS-independent (black) MM lines. Data adapted from (*13*). **K**) Excess HSA drug synergy matrices for indicated MM and AML lines treated with titrations of FRAX597 (x-axis) and RMC-6236 (y-axis). Cell viability was measured via CCK-8 at 4 days, n=3. **L**)Weights of mice from MM.1S xenograft experiment treated with indicated drugs.
